# The control of protein arginine phosphorylation facilitates proteostasis by an AAA+ chaperone protease system

**DOI:** 10.1101/2022.09.15.508104

**Authors:** Regina Alver, Ingo Hantke, Fabián A. Cornejo, Katrin Gunka, Sebastian Rämisch, Noël Molière, Emmanuelle Charpentier, Kürşad Turgay

**Affiliations:** Max Planck Unit for the Science of Pathogens, Berlin, Germany; Institut für Mikrobiologie, Leibniz-Universität Hannover, Hannover, Germany; Institut für Biologie, Humboldt-Universität zu Berlin, Berlin, Germany

**Keywords:** Stress response, protein arginine phosphorylation, protein homeostasis, chaperones, AAA+ proteins, adaptor proteins, McsB kinase, YwlE phosphatase, ClpCP

## Abstract

We could demonstrate that the AAA+ unfoldase ClpC together with the protein arginine kinase and adaptor protein McsB, its activator McsA and the phosphatase YwlE form a unique chaperone system. Here, the McsA-activated McsB phosphorylates and targets aggregated substrate proteins for extraction and unfolding by ClpC. Sub-stoichiometric amounts of the YwlE phosphatase enhanced the ClpC/McsB/McsA mediated disaggregation and facilitated the de-phosphorylation of the arginine-phosphorylated substrate protein extruded by ClpC, allowing its subsequent refolding. Interestingly, the successfully refolded protein escaped degradation by the loosely associated ClpP protease. This unique chaperone system is thereby able to disaggregate and refold aggregated proteins but can also remove severely damaged protein aggregates by degradation.

## Introduction

Molecular chaperone systems monitor and maintain cellular protein homeostasis. These include chaperone systems such as the Hsp70 & 40 or the Hsp60 system, which can refold and repair misfolded proteins. In many bacteria and yeast, AAA+ unfoldases such as Hsp104 (ClpB) form with DnaK (Hsp70) a bi-chaperone system that can disaggregate and refold and thereby remove and repair already aggregated protein species. The proteolytic arm of the bacterial protein quality control system includes AAA+ protease complexes, consisting of AAA+ unfoldases, such as ClpC, associated with a compartmentalized ClpP protease complex, allowing to degrade and thereby remove more severely damaged and possibly toxic protein aggregate species. The substrate recognition and selection by these AAA+ unfoldases often depend on adaptor proteins that can recognize and target substrate proteins. Notably, adaptor proteins, controlling the activity and stability of key transcription factors, allow AAA+ protease complexes such as ClpCP, ClpAP or ClpXP to be involved in regulatory proteolysis (*1*–*6*).

The ClpCP AAA+ protease complex of *B. subtilis* is directly involved in general and regulatory proteolysis, including heat stress response and removing subcellular protein aggregates (*7*–*11*). The substrate recognition and activation of ClpC depends on adaptor proteins, MecA, its paralog YpbH and the protein kinase McsB. The role of YpbH is not well understood (*12, 13*), MecA is involved in controlling competence development (*14*), and McsB is known for its role in controlling CtsR-mediated heat shock response (*15*).

The protein kinase activity of the McsB adaptor protein catalyzes the formation of a phospho-arginine protein modification with a high-energy phosphoramidate bond (*16*). McsB autophosphorylates itself, and in the presence of McsA, the arginine kinase activity is further induced, resulting in increased autophosphorylation and enhanced phosphorylation of McsA, ClpC and substrate proteins such as the repressor CtsR. YwlE is the cognate phospho-arginine phosphatase, which can counteract the McsB kinase activity, thereby controlling the extent of this protein modification in *B. subtilis* (*15*–*22*).

McsB, like the other adaptor proteins, can interact with the NTD of ClpC, thereby targeting substrate protein for ClpCP degradation. This activity can occur independently of the arginine phosphorylation of the substrate (*15, 23, 24*). However, a clear link was observed between the kinase activity of McsB, ClpC activation and its ability to target substrate proteins (*15, 17, 19, 22*). Recently, an additional mechanism of substrate targeting to ClpC was described, where protein arginine phosphorylation by McsB functions as a degron to mark an unfolded substrate protein for direct recognition by the N-terminal domain of ClpC, leading to subsequent degradation by ClpCP (*22, 25*).

A very pleiotropic influence of protein arginine phosphorylation in *B. subtilis* physiology and regulatory pathways, which includes *B. subtilis* spore germination (*26, 27*), was observed in a *ywlE* mutant (*17, 20*). However, the role and function of this protein modification system and the interplay of the respective kinase and phosphatase with the ClpC chaperone system in cellular protein homeostasis, is not well understood.

## Results

### McsB is the main ClpC adaptor protein allowing the *in vivo* removal of subcellular protein aggregates

To identify which of the adaptor proteins is supporting the role of ClpC in protein homeostasis, we constructed a triple adaptor mutant strain (TAM) lacking the genes for the three known ClpC adaptor proteins (Δ*mecA*Δ*ypbH*Δ*mcsB*). This TAM strain was impaired in thermoresistance and in its ability to develop thermotolerance (Fig 1A & S1A). We ectopically introduced and expressed each single adaptor protein gene in this TAM strain (Fig 1BC Fig S1BD) and observed that only the expression of *mcsB*, and not *ypbH* or *mecA*, was able to confer thermoresistance and thermotolerance at a level comparable to the wild type strain of *B. subtilis* (Fig 1B, Fig S1BC). In addition, after a 90 min recovery at 37 °C, only the expression of *mcsB* resulted in the complete removal of heat-induced sub-cellular protein aggregates, marked by the YocM-mCherry reporter (*9*), (Fig 1C, FigS1E). Furthermore, McsB-GFP co-localized *in vivo* with these subcellular protein aggregates (Fig S1FG), as observed for its substrate CtsR after a heat shock (*28*).

**Figure 1.**
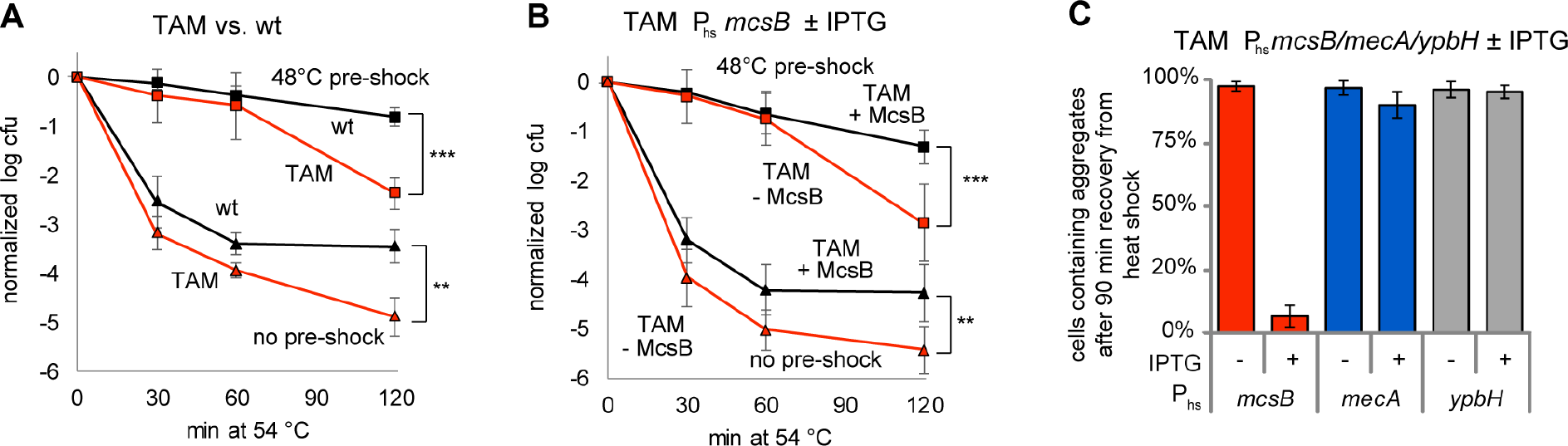
Role of the adaptor protein McsB in protein homeostasis. **A)** Thermotolerance (squares: with pre-shock) and Thermoresistance (triangles: without pre-shock) experiments analyzing *B. subtilis* Δ*mecA*Δ*ypbH*Δ*mcsB* (‘TAM’, BIH737, red) and wild-type (black) strains. **B)** *B. subtilis* TAM P_hs_ *mcsB* strain (BIH739), with (black) and without IPTG (red), were examined in a thermotolerance (squares: with pre-shock at 48 °C) and thermoresistance (triangles: no pre-shock) experiment. Log_10_-values of viable cell count were normalized, and three biological replicates were used to calculate standard deviations (error bars) with significant differences indicated (*: p<0.05, **: p<0.01, ***: p<0.001). **C**) Subcellular protein aggregates during recovery after heat stress *in vivo*. The ratio of cells containing protein aggregates marked by YocM-mCherry of complemented *B. subtilis* TAM strains (TAM P_hs_ *mscsB* (BIH739), TAM P_hs_ *mecA* (BIH740), TAM P_hs_ *ypbH* (BIH741), which were heat shocked for 30 min at 50°C and recovered for 90 min at 37°C, was determined based on three biological replicates including three technical replicates of 50 – 100 cells. Samples were taken at indicated time points for analysis by fluorescence microscopy (see also Fig S1D).

These experiments strongly suggest that McsB is the ClpC adaptor protein necessary and sufficient for thermoresistance and thermotolerance development and to target subcellular protein aggregates for ClpC-dependent removal (*9*) (Fig 1 & Fig S1).

### McsA-activated McsB kinase facilitates ClpC disaggregation but interferes with the subsequent *in vitro* refolding of aggregated Mdh

We used heat-aggregated malate dehydrogenase (Mdh) as a model substrate to reconstitute and assess the disaggregation and refolding activities of this chaperone system *in vitro* (*13, 29, 30*). In control experiments, we confirmed that the enzyme activity of folded Mdh was not significantly diminished by the active McsB kinase (Fig S1J). We could demonstrate that the well-characterized *E. coli* ClpB, DnaK, DnaJ and GrpE (ClpB/KJE) chaperone system (*29*) fully dissolved the Mdh protein aggregate and facilitated the recovery of 30% of enzyme activity (Fig S1H). Moreover, as previously observed, the ClpC/MecA chaperone system facilitated a slower disaggregation with a less effective refolding of about 15% (*13*) (Fig S1H). Adding McsB, McsA & ClpC to the heat-aggregated Mdh resulted in an apparent initial complex formation, but the subsequent disaggregation of the substrate protein was lower than observed for ClpB/KJE (Fig S1H, left panel), and only very little refolding (about 5%) could be observed (Fig S1H, right panel).

In supernatant/pellet experiments analyzed by SDS-PAGE, only the solubilization of aggregated Mdh by ClpB/KJE could be detected (Fig S1I). With an immuno-blot utilizing an antibody generated to detect protein arginine phosphorylation (*31*), we observed that most of the arginine phosphorylated proteins could be detected after 15 min, mostly in the soluble fraction (Fig S1I). This coincides with the initial disaggregation complex of ClpC/McsB/McsA appeared to become stabilized and active, as observed by light scattering (Fig S1H, Fig S1I). The relatively low refolding activity by ClpC, McsB and McsA (Fig S1H) might be explained by the unhinged McsA-induced kinase activity of McsB (Fig S1I), whereby the protein arginine phosphorylation of the substrate protein could interfere with its successful refolding.

To further assess the influence of the arginine phosphorylation by McsA and McsB on refolding, we utilized the kinase-inactive McsB C167S variant (*15, 19*), which could activate *in vitro* the ClpC ATPase activity to about 50 % of McsB-mediated activation (Fig S2C). McsB C167S could facilitate the ClpC mediated disaggregation reaction *in vitro*, albeit with a slower kinetic, but reaching a similar yield after 120 min (Fig S2E). Importantly, with McsB C167S we observed *in vitro* a significantly increased recovery of Mdh activity to about 20% of the initial activity compared to the 5% of McsB with McsA (Fig. S2E) or the 15% in the presence of MecA (Fig S1H).

While no disaggregation activity was detected *in vivo* in the absence of McsB, we observed partial disaggregation activity at about 25% when complementing a Δ*mcsB B. subtilis* strain with the McsB C167S variant compared to wild-type McsB (Fig S2B). The kinase-inactive *mcsB* C167S mutant strain was impaired by about one order of magnitude in developing thermotolerance or thermoresistance (Fig S2A). We observed a comparable *in vivo* co-localization to subcellular protein aggregates by the kinase-inactive McsB C167S-GFP and the McsB-GFP fusion proteins, suggesting that McsB can recognize subcellular protein aggregates independently of its kinase activity (Fig S2D).

In summary, the kinase-inactive McsB C167S variant behaved to a certain extent like a ClpC adaptor protein and allowed, in contrast to kinase-active McsB, a significant refolding of the unfolded substrate protein (Fig S2E & S2F), suggesting that active substrate phosphorylation by McsB might interfere with the subsequent refolding reaction.

### The presence of sub-stoichiometric amounts of the YwlE phosphatase facilitates McsB kinase-mediated *in vitro* disaggregation and refolding by ClpC

These experiments suggested that controlling the protein arginine phosphorylation state might be very important for the disaggregation and refolding activity of the McsB/McsA/ClpC chaperone complex (*17, 20*). Previous *in vitro* experiments demonstrated that equimolar amounts of YwlE can inhibit the McsB and McsA-mediated activation of ClpC (*15*). However, when we lowered the YwlE concentration, we observed that at sub-stoichiometric levels of YwlE, the McsA & McsB-induced ClpC ATPase activity was no longer inhibited (Fig 2A).

**Figure 2.**
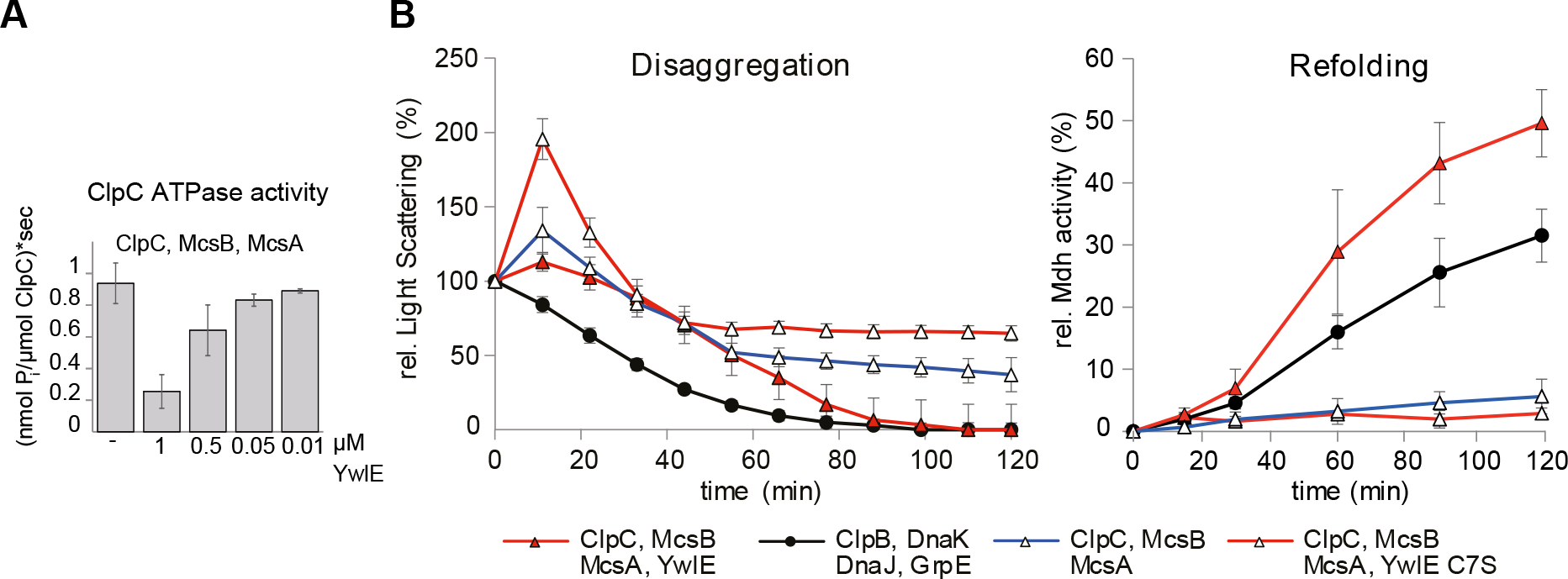
Sub-stoichiometric YwlE facilitates *in vitro* disaggregation and refolding. **A**) ATPase activity of ClpC/McsB/McsA (1μM) was determined in the absence or presence of indicated amounts of YwlE (0.01-1μM). **B**) Disaggregation and refolding experiments. Disaggregation of heat-aggregated Mdh after the addition of the indicated proteins was monitored *in vitro* by measuring light scattering at 30°C for 120 min (left panel). The refolding was determined by Mdh enzymatic activity (right panel). 0.05 μM YwlE or YwlE C7S were added when indicated. The plots show the mean of at least three replicates with standard deviation.

We added this sub-stoichiometric amount of YwlE (0.05 μM) to the *in vitro* disaggregation and refolding assay with ClpC, McsB & McsA (at 1μM) and observed that about 100% of the heat-aggregated Mdh was disaggregated before 90 min, and 50% was successfully refolded after 120 min (Fig 2B). Accordingly, pellet-supernatant experiments displayed a clear shift of Mdh from the pellet to the soluble fraction in the presence of ClpC, McsB & McsA with sub-stoichiometric amounts of YwlE (Fig S3A).

Interestingly, arginine protein phosphorylation was close to or even below the detection level by western blot analysis, suggesting that sub-stoichiometric amounts of active YwlE are sufficient to lower the phosphorylation state of the involved proteins effectively (Fig S3A).

The addition of the phosphatase inactive YwlE C7S variant (*18*) to the disaggregation and refolding experiment resulted in a low disaggregation rate without apparent refolding (Fig 2B). Consequently, no solubilization of aggregated protein could be observed in the respective supernatant pellet experiment with YwlE C7S (Fig S3A). In the presence of the YwlE C7S variant, the concomitant protein arginine phosphorylation was clearly detectable with a maximum after 15 min, like in the absence of YwlE (Fig 2B & S3A). With order-of-addition experiments, we observed that the later the addition of YwlE was, the lower the disaggregation and the refolding yields were (Fig S4AB).

These experiments suggest that a specific YwlE phosphatase activity is necessary for efficient disaggregation and especially refolding by the ClpC/McsB/McsA chaperone system. YwlE-mediated de-phosphorylation might directly support refolding of unfolded substrate proteins (Fig 2B) and could also modulate ClpC, McsB and McsA activities (*15, 17, 20, 32*) (see Fig 6A).

### A well-balanced YwlE level is necessary for functioning *in vivo* protein homeostasis

The sub-stoichiometric levels of YwlE relative to the ClpC chaperone complex from the *in vitro* experiment of about 1:20-1:30 (Fig 2) correspond relatively well to the observed *in vivo* ratio of about 20 to 100 times lower cellular concentration of YwlE, compared to the higher abundant ClpC determined *in vivo* for *B. subtilis* cells by different methods in various conditions (*14, 33, 34*).

Since the relative levels of the YwlE phosphatase appear to be important for the *in vitro* disaggregation and refolding, we went on to investigate the effect of different cellular *in vivo* levels of YwlE on protein homeostasis and heat shock response. For this, we used the Δ*ywlE* mutant strains where we could control cellular YwlE levels with an IPTG inducible expression of *ywlE* or *ywlE* C7S *in trans*. We measured the cellular YwlE levels of wild-type *B. subtilis* and these strains by quantitative western blotting and observed about 134 (±34) YwlE molecules per cell in wild-type and 64 (±11) in the complemented Δ*ywlE* P_hs_ *ywlE B. subtilis* strain without induction (Fig 3D & Fig S3E). The addition of 10 μM IPTG doubled the cellular YwlE to 274 (± 26) molecules/cell and reached about 6500 (± 1702) YwlE molecules per cell when 2 mM IPTG was added (Fig 3D & S3E). Interestingly, the uninduced Δ*ywlE* P_hs_ *ywlE* strain with almost wild-type levels of YwlE behaved like the wild-type in thermotolerance and thermoresistance experiments. The more IPTG inducer was added, the more YwlE was expressed and the less thermoresistant or thermotolerant these cells became (Fig 3AC & S3C). However, thermotolerance and thermoresistance were as low as in a Δ*ywlE* strain when complemented with the phosphatase inactive *ywlE* C7S variant (Fig 3C). Assessing the subcellular aggregate formation revealed increased subcellular aggregates at 48°C and 48/53°C during thermotolerance for the cells with increased *ywlE* expression but not when *ywlE* C7S was expressed (Fig S3B). Furthermore, high levels of *ywlE* diminished the clearance of heat-induced aggregates during recovery from heat stress (Fig 2D), which was not observed with low YwlE levels or for the Δ*ywlE* strain complemented with the highly expressed *ywlE* C7S (Fig 3B).

**Figure 3.**
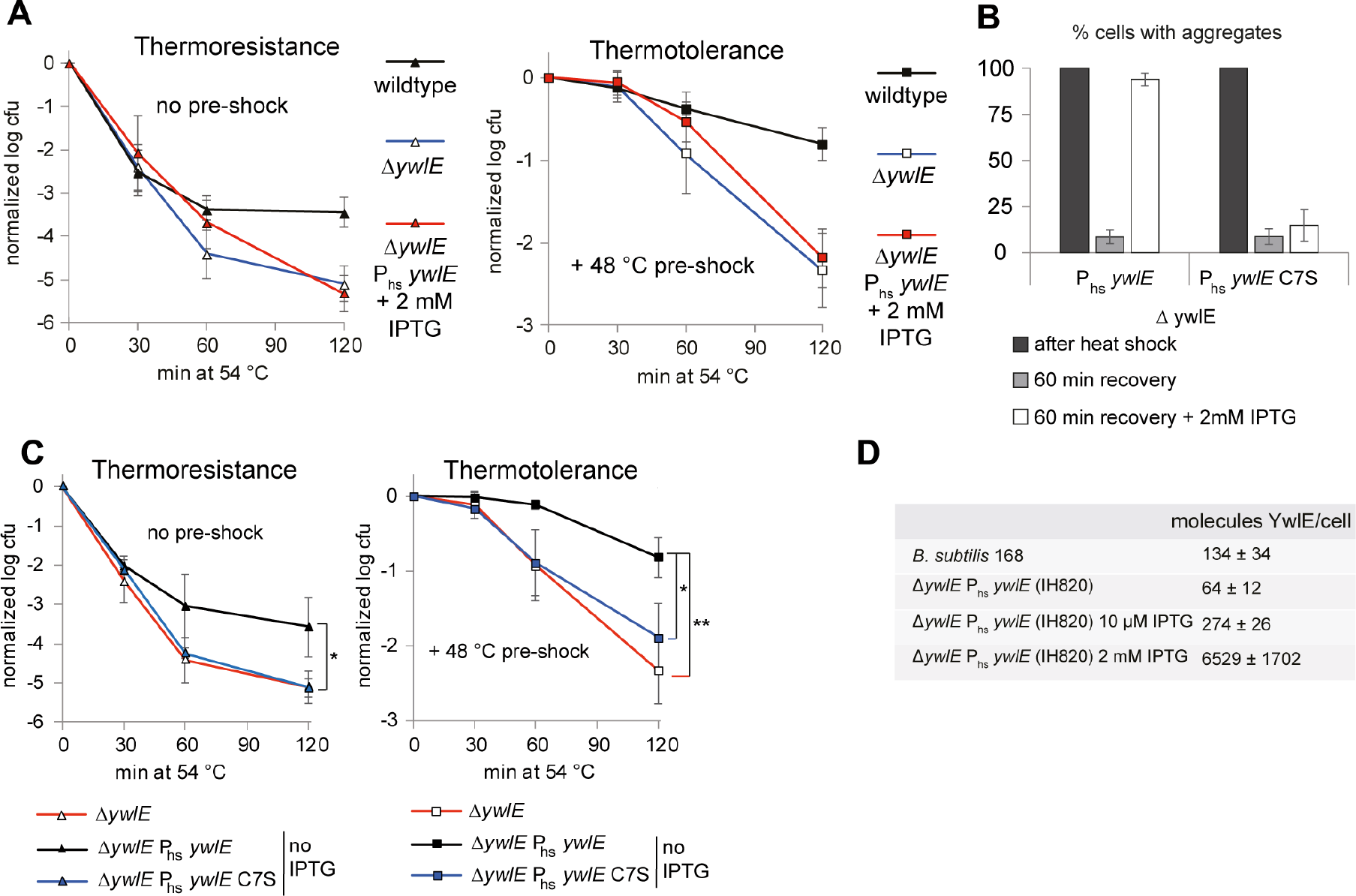
Low *in vivo* levels of YwlE are necessary to facilitate protein homeostasis. **A**) Thermoresistance and thermotolerance experiments with *B. subtilis* wild-type (black), Δ*ywlE* (BIH309, blue) and Δ*ywlE* P_hs_ *ywlE* + 2 mM IPTG (BIH828, red) were performed. Log_10_-values of viable cell counts were normalized, and three biological replicates were used to calculate standard deviations (error bars). **B**) Subcellular protein aggregates during recovery after heat stress *in vivo*. Cells of *B. subtilis* strains Δ*ywlE* P_hs_*ywlE* (BIH828) and Δ*ywlE* P_hs_ *ywlE* C7S (BIH829) both also encoding P_xy_*yocM-mCherry* were heat shocked for 20 min at 52°C and recovered for 60 min at 37°C. Cells were taken at the indicated time points, and their subcellular protein aggregates analyzed by fluorescence microscopy. Three biological replicates, including three technical replicates with about 50 – 100 cells were analyzed to calculate the ratio of cells containing protein aggregates marked by YocM-mCherry. **C**) Thermoresistance and thermotolerance experiments with *B. subtilis* Δ*ywlE* (BIH309, red), Δ*ywlE* P_hs_*ywlE* (BIH828, black) and Δ*ywlE* P_hs_ *ywlE* C7S (BIH829, blue) without IPTG. Log_10_-values of viable cell count were normalized, and three biological replicates were used to calculate standard deviations (error bars) with significant differences indicated (*: p<0.05, **: p<0.01, ***: p<0.001, Welch’s test). **D**) Table showing the number of YwlE proteins/cell determined *in vivo* by quantitative Western blotting for the different *B. subtilis* strains at indicated IPTG concentrations (see Fig S3E).

We did not observe an influence of *ywlE* expression on the level of subcellular aggregates during recovery from heat stress in a kinase-inactive *mcsB*C167S mutant (Fig S3D), confirming that the presence of protein arginine phosphorylation activity is required to observe an influence of the counteracting *ywlE* phosphatase activity.

High levels of YwlE phosphatase interfered strongly with the McsB kinase and thereby appeared to behave like a strain lacking McsB (Fig S2AB & Fig3AC) both *in vitro* and *in vivo*. Controlling the cellular levels of protein arginine phosphorylation is therefore not only mediated by the McsA-activated McsB kinase but also controlled by the YwlE phosphatase, facilitating a well-balanced equilibrium of cellular protein arginine phosphorylation levels to maintain cellular protein homeostasis.

### The YwlE-dependent *in vitro* refolding is independent of the ClpC unfolding activity

To assess the interplay of the ClpC chaperone activity and the refolding facilitated by the YwlE phosphatase, we used a ClpC trap variant (ClpC-DWB), which cannot hydrolyze ATP. ClpC-DWB can form mixed oligomers with ClpC and can thereby interfere with the ATPase driven activities of ClpC (Fig S4C & S4D) (*35*–*38*). The addition of ClpC-DWB to our *in vitro* assay with ClpC, McsA & McsB 15 min after the start, resulted in a slow decline to a lower but constant light scattering signal for the remaining time (Fig S4C), suggesting that the disaggregation was halted (*36*). The simultaneous or subsequent addition of YwlE (at 15 (green squares) or 20 min (blue diamonds)) appeared to disrupt the apparent complex much faster, resulting in a drop to the same lower level of constant light scattering (Fig S4C). Importantly, the addition of active YwlE could facilitate up to 15 % recovery of enzyme activity after 120 min (green squares). However, no considerable refolding occurred when YwlE C7S variant was added (purple circles) (Fig S4C). These experiments suggest that the YwlE phosphatase activity is necessary to support the refolding of the available unfolded and phosphorylated substrate proteins independent of a concurrent ClpC ATPase-driven unfoldase activity (Fig S4C).

### McsA is necessary for starting the disaggregation and refolding activity

We could demonstrate that adding McsA at different time points to McsB, ClpC and YwlE immediately increased the light scattering signal suggesting an initial protein complex formation, followed by the onset of the disaggregation reaction and Mdh activity starting to become recovered (Fig S4E). At the same time, the start of protein arginine phosphorylation of the involved proteins could be detected, confirming the concurrent activation of the McsB kinase activity by McsA (Fig S4F) (*19*). These experiments suggest that the McsA interaction with McsB and its kinase activation is necessary to initiate the efficient disaggregation and subsequent refolding activity by this chaperone system.

### ClpP association with ClpC: allowing a switch between repair or removal?

ClpC and ClpP can form the ClpCP AAA+ protease complex (*39*–*42*). Therefore, we investigated how the presence of ClpP influences the disaggregation and refolding activity of McsB, McsA, ClpC, and YwlE (Fig 4A) and to what extent this repair activity would shift toward substrate degradation (Fig 4B).

**Figure 4.**
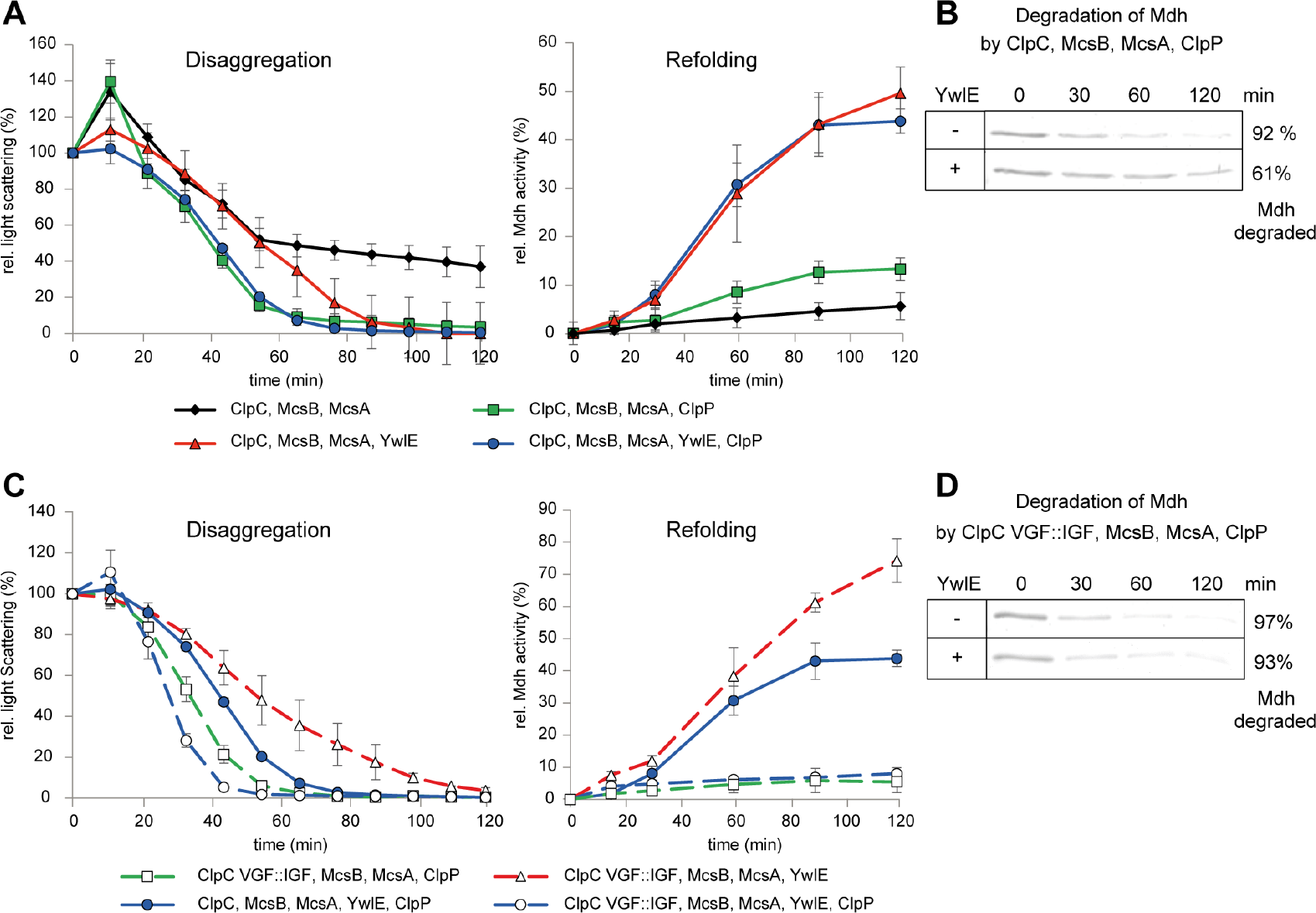
ClpCP interaction strength and the influence of YwlE on refolding or degradation. Disaggregation, refolding and degradation experiments. **A**) After the addition of the indicated proteins, disaggregation of heat-aggregated Mdh (1 μM) was monitored *in vitro* by measuring light scattering (left panel) and refolding by measuring the Mdh enzymatic activity (right panel). **B**) *In vitro* degradation experiments of heat aggregated Mdh as substrate were performed with the indicated proteins. Samples were taken for analysis by SDS PAGE and Coomassie stain. The relative amount of degraded Mdh after 120 min is indicated. **C**) Disaggregation of heat aggregated Mdh (1 μM) was monitored *in vitro* after the addition of the indicated proteins by measuring light scattering (left panel) and refolding by measuring the Mdh enzymatic activity (right panel) **D**) *In vitro* degradation experiments of heat-aggregated Mdh as substrate were performed with the indicated proteins. Samples were taken for analysis by SDS PAGE and Coomassie stain. The relative amount of degraded Mdh after 120 min is indicated.

Adding ClpP in the absence of YwlE significantly enhanced the disaggregation activity (Fig 4A, green squares), and about 90% of aggregated Mdh was degraded (Fig 4B). Still, about 10% of the aggregated Mdh was refolded (Fig 4A, green squares). The raised disaggregation activity (Fig 4A) is consistent with the observed concurrent induction of the ClpC ATPase activity when ClpP was added (Fig S5A). The additional presence of YwlE did not further influence disaggregation efficiency but resulted in about 40 % refolding of the aggregated Mdh (Fig 4A, blue circles), similar to the reaction in the presence of YwlE but without ClpP (Fig 4A, red triangles). At the same time, about 60% of the initial aggregated Mdh (Fig 4A, blue circles) was degraded (Fig 4B). These *in vitro* experiments suggest a triage where the refolded Mdh species could escape the transfer to the associated ClpP protease, but the still unfolded substrate proteins were successfully transferred for degradation into the ClpP protease complex.

The ClpC association with ClpP is mediated by the tips of ClpP-interacting loops (P-loops) extending from the bottom of the AAA+ protein hexamer and interacting with binding pockets on the apical surface of the double-heptameric ClpP complex (*6, 39, 40, 42, 43*). The P-loop of ClpC contains a VGF motif at its tip, while other Clp proteins like ClpX from *B. subtilis* and *E. coli* possess IGF P-loops to facilitate this interaction.

Interestingly, it was demonstrated that in the context of *E. coli* ClpXP, the ClpX IGF loop allows a tighter interaction between ClpX and ClpP than a VGF loop (*39*). To further evaluate the difference between the VGF vs IGF P-loop for the interaction of ClpC with ClpP, we utilized the recently resolved structure of a ClpX-ClpP complex from *Listeria monocytogenes* (*44*) and the *B. subtilis* ClpP structure (*45*), to model and calculate thermodynamic interaction parameters of the IGF compared to the VGF P-loop for ClpP binding. These *in silico* modeling and calculations suggested a stronger binding interaction of the IGF P-loop than the VGF P-loop (Fig S5E). When we replaced the Valine in the VGF motif with Isoleucine, we observed in our *in vitro* experiments that the ClpC VGF::IGF variant in the presence of McsB and McsA displayed a slightly lower ATPase (Fig S5B) and a lower disaggregation activity (Fig 4C), and a high refolding activity of up to 80% in the presence of YwlE (Fig 4C). Adding ClpP to ClpC VGF::IGF with McsB and McsA enhanced the disaggregation activity and switched the proteolytic activity of this complex to a full degradation of Mdh, independent of the presence of YwlE (Fig 4D, Fig S5D). The observed complete degradation of the substrate protein by the ClpC VGF::IGF/ClpP AAA+ protease complex (Fig 4D) contrasts the only partial degradation of the remaining unfolded substrate protein with wild-type ClpC (Fig 4B). This difference suggests that a closer and possibly tighter interaction of ClpC VGF::IGF with ClpP might interfere with the escape of already folded substrate protein observed for wild-type ClpC-ClpP (Fig 4AB).

Such a relatively transient, more dynamic ClpC-ClpP interaction during disaggregation and unfolding would give better access to the YwlE phosphatase for the de-phosphorylation of translocated phosphorylated unfolded protein substrates. And it could allow at the same time, a more unconstrained folding of the extruding nascent polypeptide chains (see Fig 6B).

### The ClpCP AAA+ protease complex formation is not necessary for protein homeostasis during heat shock but beneficial during more severe proteotoxic stress

Our *in vitro* experiments suggest that the ClpCP complex can switch between a refolding and a degradation mode, depending on the interaction strength of its P-loop with ClpP (Fig 4). To investigate the role of ClpC/ClpP interaction *in vivo*, we constructed a *B. subtilis* mutant strain, encoding a P-loop ClpC VGF::GGR variant, which abolishes the interaction with the ClpP complex and has, therefore, no proteolytic activity towards known substrates neither *in vivo* nor *in vitro* (Fig S6AB) (*14, 15*). However, it is active in the *in vitro* disaggregation and refolding experiment (Fig S6E), with a slightly lower ATPase compared to ClpC (Fig S6D).

While a Δ*clpC* deletion mutant showed impaired thermotolerance development, less thermoresistance and increased protein aggregation during heat shock (*9*–*11*), a *clpC* VGF::GGR and the *clpC* VGF::IGF mutant strain behaved comparable to the wild-type strain in thermoresistance, thermotolerance (Fig 5A & Fig S5C) and protein aggregate formation (Fig S6C). In addition, McsB-mediated removal of heat-induced protein aggregates during recovery from heat shock was not affected in a wild-type or a *clpC* VGF::GGR, but strongly impaired in the *clpC* DWB background strain (Fig 5BC). Interestingly, fewer subcellular protein aggregates were detected in the *clpC* VGF::IGF mutant strain already without induction of McsB expression. However, upon full McsB induction, they were removed to the same level as in the wild-type ClpC control (Fig 5C).

**Figure 5.**
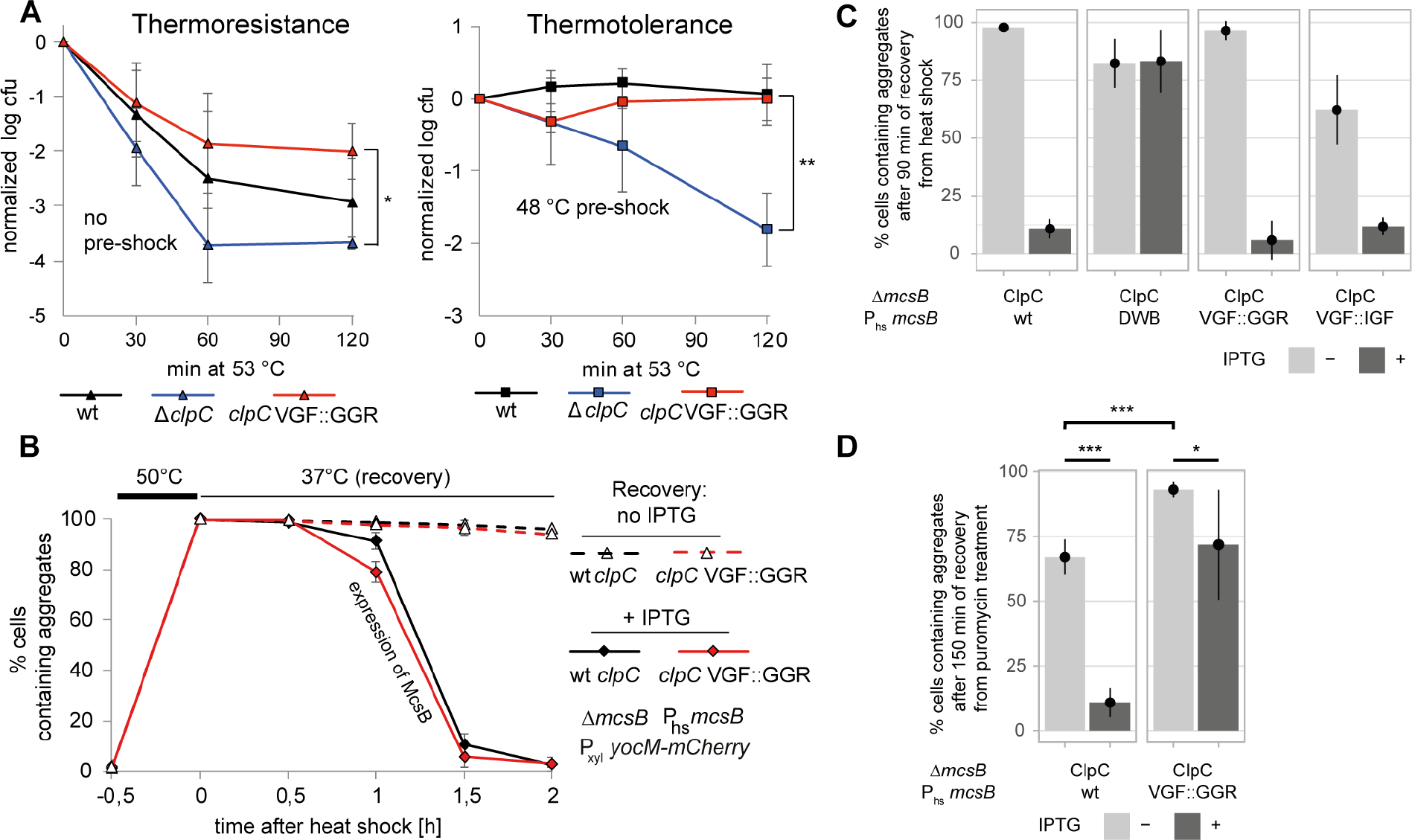
ClpC P-loop variants: rescue or removal? **A**) Thermoresistance and thermotolerance experiments with *B. subtilis* wild-type (black), Δ*clpC* (BIH19, blue) and *clpC* VGF::GGR (BIH217, red) cells. Log_10_-values of viable cell count were normalized, and three biological replicates were used to calculate standard deviations (error bars). Significant differences are indicated (*: p<0.05, **: p<0.01, ***: p<0.001, Welch’s test). **B**) Cells of *B. subtilis* ClpC wt (BIH414, black) and ClpC VGF::GGR (BIH485, red) (with Δ*mcsB* P_hs_ *mcsB* P_xyl_ *yocM-mCherry*), were treated with a 50°C heat shock and subsequently shifted to recover at 37°C. The ratio of cells containing protein aggregates were determined. **C**) ClpC wt (BIH414), ClpC-DWB (BIH488), ClpC VGF::GGR BIH485) and ClpC VGF::IGF (BIH860) strains (and Δ*mcsB* P_hs_ *mcsB* P_xyl_ *yocM-mCherry*) were treated with a 50°C heat shock and shifted for 90 min to 37°C in the absence or presence of IPTG. The ratio of cells containing protein aggregates was determined. Three biological replicates, including three technical replicates with 50 – 100 cells, were analyzed. Log_10_-values of viable cell count were normalized, and three biological replicates were used to calculate standard deviations (error bars). **D**) *B. subtilis* ClpC wt (BIH414) and ClpC VGF::GGR (BIH485) were treated with 25 μg/mL puromycin. After 15 min, puromycin was removed, and the cells were incubated in fresh media with xylose in the presence or absence of 2 mM IPTG at 37 °C for 150 min. Samples were taken for analysis by fluorescence microscopy and western blot analysis (see also S6F & G). The ratio of cells containing protein aggregates was determined based on three biological replicates, including three technical replicates with 50 – 100 cells. Log_10_-values of viable cell count were normalized and three biological replicates were used to calculate standard deviations (shown as error bars) with significant differences indicated (*: p<0.05, **: p<0.01, ***: p<0.001, Welch’s test).

We also applied a more severe protein folding stress by adding the antibiotic puromycin, a tRNA mimic whose presence results in premature translation termination, generating easily aggregating non-functional protein fragments (Fig 5D & Fig S6F). After removing puromycin, we observed the clearance of protein aggregates in the *mcsB* deletion mutant after 150 min, only when *mcsB* expression was induced (Fig 5D & Fig S6F). However, the clearance of these truncated protein aggregates was strongly diminished in the *clpC* VGF::GGR mutant (Fig 5D & Fig S6F).

These experiments demonstrated that the ClpC VGF::GGR variant, although not able to associate with ClpP, could very well replace wt ClpC as unfoldase, also during heat shock response, however, not when more toxic not refoldable protein aggregates were present (Fig 5ABC, Fig S6C). This suggests that switching to ClpCP-mediated degradation can be beneficial under more severe protein folding stress (Fig 6).

**Fig 6.**
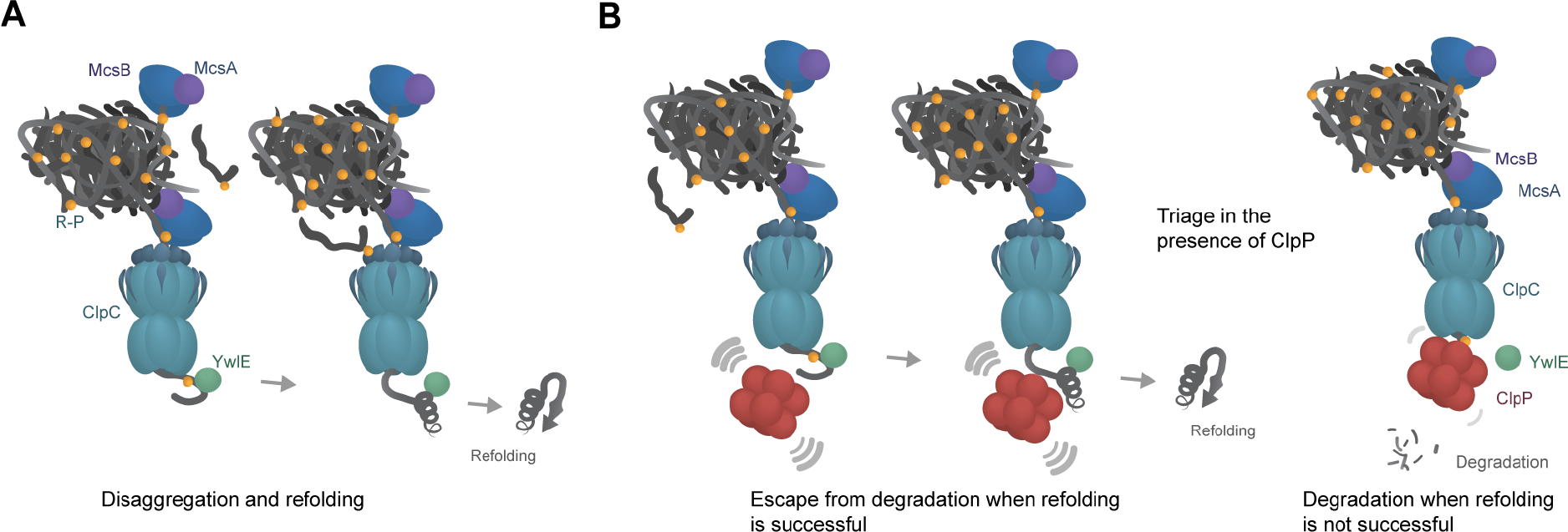
Disaggregation and refolding or degradation. **A)** Model for the disaggregation and refolding in the absence of ClpP. **B)** Model for the triage in the presence of ClpP: Escape from transfer to degradation by extruded and simultaneously refolded protein substrates (left). Degradation of not folded protein substrates.

## Discussion

Here, we could identify and describe an unusual AAA+ chaperone system, which integrates protein modification and de-modification to enhance and facilitate cellular protein homeostasis. In this chaperone system, the cellular level of protein arginine phosphorylation is controlled by sub-stoichiometric level of the YwlE phosphatase. Thereby the McsB kinase activity and cellular protein arginine phosphorylation can be limited, which allows the modulation of ClpC-mediated disaggregation and unfolding activities and the additional subsequent facilitation of substrate refolding (Fig 2 & 3). The specific targeting of misfolded or aggregated protein by McsB (*28*) (Fig 1, 2, & S2) is consistent with the previous identification of a wide array of cellular proteins, which can become modified by protein arginine phosphorylation, since they might be prone to aggregation (*17, 20*).

The kinase activity of the McsB adaptor protein adds complexity but also expands the recognition and targeting abilities of McsB. The activation of ClpC by McsB can be facilitated by binding to the NTD of ClpC, but ClpC can be further activated by phosphorylation of specific arginine residues located in the NTD of ClpC (*17, 20, 22*). We could observe a basal adaptor protein activity for the kinase-inactive McsB C167S variant (Fig S2). Both McsB and McsB C167S interact with the NTD of ClpC, and both recognize sub-cellular protein aggregates (Fig S2) and can directly target misfolded or aggregated substrate proteins to ClpC, even when the substrate proteins lack arginines (*15, 17, 24*). This ability of McsB would also allow the recruitment of ClpC to these substrate protein structures (*15, 28*), similar to DnaK, which can act as an adaptor protein for the ClpB disaggregase (*46*).

A distinct substrate recognition and targeting pathway involving the McsB kinase was reported, where McsB is not necessarily interacting directly with ClpC. Here the phospho-arginine modifications of unfolded substrate proteins, generated directly by the McsB kinase, can act as a degron, which allows the direct recognition of the marked substrate proteins via specific phospho-arginine interaction sites in the NTD of ClpC (*21, 22, 25*). This additional second recognition path for soluble substrate could play a role for a more efficient function of the chaperone complex since unfolded phosphorylated substrate proteins that exit ClpC and are neither dephosphorylated by YwlE, nor degraded by ClpP could then become directly retargeted by their still present degron to ClpC unfolding (*22*) (Fig 6).

The arginine phosphorylation of unfolded, misfolded, or aggregated protein species might change the physical properties of these protein species in different ways. Arginine phosphorylation reverts a positive to a negative charge, which can result in obliterating unspecific or specific interaction with DNA or RNA (*16*). Protein aggregates with these differently charged misfolded phosphorylated protein species might be physically better accessible. And ClpC could also interact directly via its phospho-arginine receptors of the NTD with protein arginine phosphorylated misfolded proteins exposed on the surface or in the vicinity of such protein aggregates (*22*).

We observed that the YwlE phosphatase was essential to facilitate the refolding of the unfolded phosphorylated protein substrates extruded from the ClpC chaperone system (Fig 2 & 4). Here we hypothesize that the released unfolded and phosphorylated substrate protein species might be unable to refold successfully. However, in the presence of YwlE, the phosphoryl group can be removed, possibly allowing the refolding of the substrate protein, which is somehow reminiscent of the classical refolding experiments by Anfinsen (*47*).

Notably, a high-energy P-N bond is formed in phospho-arginine by McsB (*16, 48*). This might be an important aspect when considering possible thermodynamic requirements of the folding process accompanying the YwlE-mediated protein arginine de-phosphorylation.

Different combinations of specific Hsp40, Hsp70, and Hsp110 chaperone systems or AAA+ unfoldases, such as ClpB (Hsp104) combined with Hsp70, were demonstrated to remove protein aggregates by disaggregation and refolding in their respective cellular systems. Some AAA+ unfoldases, such as ClpG, even displayed a stand-alone disaggregation and refolding activity (*5, 49, 50*).

Here the ClpC AAA+ chaperone system supported by adaptor proteins such as MecA (Fig S1) or the kinase-inactive McsB C167S with McsA (Fig S2), could, to a certain extent, facilitate a disaggregation and refolding activity. However, coupling the ClpC-mediated unfolding to protein arginine phosphorylation and de-phosphorylation by McsB, McsA & YwlE resulted in an improved disaggregation and a significantly increased refolding activity (Fig 2). This suggests a synergy between the ClpC-mediated generation of unfolded extruding polypeptide chains, its concurrent McsB-mediated modification and the subsequent YwlE-catalyzed de-modification.

Protein stability, folding and function are known to be intricately connected to post-translational protein modification (*51, 52*). It is well established that protein modification by glycosylation and de-glycosylation play an important role in protein homeostasis in the endoplasmic reticulum (*53*–*55*). Interestingly, ubiquitination and de-ubiquitination by the deubiquitinating enzyme USP19 were suggested to support membrane protein folding at the cytosolic side of the ER membrane (*56*).

In the presence of ClpC-associated ClpP, unfolded or misfolded substrate proteins are targeted for ClpP degradation. However, already folded protein species are not transferred for degradation, due to the dynamic interaction of the P-loops with the ClpP oligomer (Fig 4). Such a relatively loose ClpC-ClpP interaction could allow YwlE access to the egressing unfolded peptide from ClpC to facilitate refolding by removing the possibly interfering arginine modification. This suggests that the McsB/McsA/ClpCP/YwlE complex can switch between repair or degradation, depending on the folding kinetics or state of the modified or already demodified nascent polypeptide chain unfolded and translocated by ClpC (Fig 6).

In the presence of the YwlE phosphatase, ClpC, McsB, and McsA establish a unique and effective disaggregation and refolding system, employing reversible protein arginine modification for various tasks, including substrate protein refolding. This ability to remove or repair stress-induced cellular protein aggregates allows to maintain protein homeostasis and is giving the bacterial cells the means to respond and withstand protein misfolding stress.

## Materials & Methods

### General methods

*B. subtilis* strains were grown in Lysogeny Broth medium (5 g l^-1^ yeast extract, 10 g l^-1^ tryptone-peptone, 5 g l^-1^ NaCl) at 37°C. Supplemental Table S1 shows the strains derived from transformations of indicated plasmids or PCR products (Table S2) derived from cloning with primers/restriction enzymes (Table S3) with standard cloning procedures (restriction enzymes and T4 DNA ligase from NEB) as described previously (*57*). Cloning was performed in *E. coli* DH5α (Invitrogen) (ampicillin 100 μg ml^-1^, kanamycin 50 μg ml^-1^) with Phusion High Fidelity Polymerase (NEB). Transformation into *B. subtilis* with chromosomal or plasmid DNA was performed by standard methods (*58*). Successful *amyE* insertion in *B. subtilis* was identified by assessing α-amylase activity on plates with 0.4 % starch (w/v) and Lugol’s iodine. Insertions in *amyE* and other genomic sites such as *lacA* were confirmed by PCR.

### Protein purification

Protein production was performed in *E. coli* BL21 (DE3) cells. Cells were grown at 37°C to an OD_600_ of 0.7 and induced with 1 mM IPTG overnight at 16°C. Purification of His_6_-tagged proteins was performed by nickel affinity chromatography after cell lysis by French press. YocM, McsB and McsB C167S were expressed and purified as N-terminal His-SUMO fusion proteins, which were released by overnight cleavage with Ulp-protease at 4°C (Andréasson et al., 2008). ClpC, ClpC-DWB, ClpC-VGF:IGF, ClpC-VGF:GGR, ClpP, MecA, McsA, YwlE and YwlEC7S were purified as previously described (*19, 36*).

### Thermotolerance development and thermoresistance

Thermotolerance development and thermoresistance experiments were performed as previously described (*9, 59*). Cell cultures were separated at OD_600_ 0.4 and treated ± a 15 min pre-shock of 48°C before both samples were shifted to 53°C or 54°C. Samples were taken for microscopy or plated on LB agar at t_0_, t_30_, t_60_ and t_120 min_. Obtained *cfu* were plotted against time. At least three biological replicates were used to calculate standard deviations, and statistical significance was calculated using Welch’s test.

### Fluorescence microscopy

*B. subtilis* cells were inoculated with a fresh overnight culture to an OD_600_ of 0.05 in LB + 0.5 % xylose (w/v) to express *yocM*-*mCherry* fusion (*9*). Fluorescence microscopy was performed at indicated time points on an object slide coated with 1 % agarose (w/v) in saline (0.9 % NaCl (w/v)). Oil immersion and 10×100 magnifications were used with a Zeiss fluorescence microscope with GFP and mCherry filters.

### *In vivo* disaggregation assay

*B. subtilis* strains grown in LB + 0.5 % xylose (w/v) were treated at OD_600_ 0.25 – 0.3 with a heat shock at 52°C for 20 min or 50°C for 30 mins if the strain lacks *mcsB*, which renders them more sensitive towards heat stress (Fig 1C). Cultures were separated and ± 2 mM IPTG was added while shifting the cells to 37°C. Samples were taken for plating (colony forming units (CFU)), western blotting and/or fluorescence microscopy as indicated. At least three biological replicates were used to calculate standard deviations, and statistical significance was calculated using Welch’s test.

### Biochemical assays

Light scattering and Mdh activity assays: Malate dehydrogenase (Roche) was inactivated and aggregated by incubating 2 μM for 30 min at 47°C. Light scattering was determined at 360 nm excitation and emission wavelength with a FP-6500-Spectrofluorometer (Jasco) at 30 °C. 1.5 μM ClpB, 1 μM DnaK, 0.2 μM DnaJ, 0.1 μM GrpE, 1.5 μM ClpC, 1 μM ClpP, 1 μM MecA, 1 μM McsB, 1 μM McsA, 0.05 μM YwlE, proteins were incubated as indicated in the respective experiments with 1 μM heat-treated Mdh in refolding buffer (50 mM Tris pH 7.8, 150 mM KCl, 20 mM MgCl_2_, 2 mM DTT) supplemented with 20 μg/ml pyruvate kinase, 4 mM phosphoenolpyruvate and 6 mM ATP as indicated in the respective experiments.

Mdh enzyme activity was determined before and after inactivation and at indicated time points by diluting 2.5 μl reaction mixture in 125 μl Mdh activity buffer (150 mM potassium phosphate buffer pH 7.6, 1 mM DTT, 0.5 mM oxalacetate, 0.28 mM NADH). The time-dependent NADH oxidation by Mdh was monitored at 340 nm in a Spectra Max MP3 Microplatereader (Molecular Devices). At least three replicates of these experiments were analyzed, and the resulting standard deviation was included as error bars.

We also followed the solubilization of aggregated proteins by supernatant pellet experiments. Samples of the disaggregation and refolding experiments were taken at indicated time points, and centrifuged for 30 min, 14000 rpm at 4°C. The supernatant was carefully separated from the pellet and mixed with the appropriate amount of 2x SDS sample buffer while the pellet was dissolved in 1x SDS sample buffer to the same final volume. Both samples were incubated for 10 min at 90°C and analyzed via SDS-PAGE and Coomassie stain (Schlothauer et al., 2003). The in vitro degradation of aggregated Mdh or β-casein in the presence of ClpP was analyzed by SDS-PAGE and Coomassie stain. The relative amount of Coomassie-stained protein band was calculated using ImageJ analysis software to determine band intensities (*60*).

The ClpC ATPase activity was measured by phosphate release, colorimetrically detected with the ammonium molybdate/malachite green assay as described previously (*61, 62*). For this assay we used 1 μM ClpC, 1 μM ClpP, 1 μM MecA, 1 μM McsB, 1 μM McsA, 0.05 μM YwlE, and possible substrates such as 1 μM Mdh, 1 μM β-casein, as indicated and if not mentioned otherwise. At least three replicates of the ATPase experiments were implemented, and the resulting standard deviation was included as error bars.

### Western blotting

SDS PAGE (15 % gels) loaded with similar amounts of protein (e.g. 5 μg total protein for cell extracts) was performed with standard methods as previously described (Laemmli 1970) with anti-YwlE, anti-ClpC, anti-MecA, anti-McsB or anti-YpbH (Pineda Berlin) antiserum as primary antibody and anti-rabbit alkaline phosphatase or horseradish peroxidase conjugate as secondary antibody (1:10000). Blots were developed with standard BCIP (5 % (w/v) in DMF) + 250 μL NBT (5 % (w/v) in 70 % DMF) or using the ECL western blotting reagent (GE Healthcare).

For phospho-arginine protein detection, samples were taken from light scattering assays without heat treatment prior to loading. SDS-PAGE and western blotting on nitrocellulose were performed at 4 °C. The primary anti-phosphoarginine-specific antibody was used 1:3000 with 5 % skim milk powder in TBS. Secondary antibodies and imaging was performed as described above (*31*). hs

### YwlE quantification

*B. subtilis* 168 (wild-type) and IH820 (Δ*ywlE*::*kanR* P_hs_ *ywlE* spec) cells were grown to an OD 0.35 ± 0.02. Subsequently, IPTG was added to a final concentration of 0.01 and 2 mM IPTG to the IH820 cultures as indicated, which were further incubated for 30 min (Fig S3C). To determine the number of bacteria, an aliquot was fixed with 3.7% formaldehyde in PBS and cells were counted in a Petroff-Hauser chamber. Cell extracts were prepared by sonication and western blotting was performed as mentioned above with few modifications. Samples were resolved in 4-15% Mini-PROTEAN TGX precast gels and transferred to 0.22 μm PVDF membranes. Anti-YwlE serum was used in a 1:5,000 dilution in 5% skim milk in TBS-T, and anti-rabbit IgG antibodies from goat conjugated to HRP were used 1:20,000 in the same buffer. Blots were developed using a chemiluminescent substrate (Thermo Scientific, 34577) and documented in a Fusion FX imaging system (Vilber). Defined amounts of purified YwlE-His were included as a calibration curve on each gel (Fig S3C). The concentration of purified YwlE-His was quantified by MS. Briefly, defined amounts of synthesized heavy isotope-coded YwlE peptides (Table S4; JPT Peptide Technologies, Berlin) were spiked into a sample of the purified YwlE protein. The concentration of the YwlE protein was then determined by dividing the MS signal of the endogenous “light” peptides by the synthetic “heavy” peptides. Western blot images were processed using Fiji (*63*).

### Structure modelling

As a structural template for modelling the ClpC-ClpP interactions, we used the ClpP-ClpX complex structure (PDB IDs: 6SFW, 6SFX) (*44*). Each ClpX subunit and the two interacting ClpP subunits were extracted, and energy minimization was performed using RosettaScripts (*64*) with the FastRelax (*65*) mover and strong coordinate constraints. To compare binding energies, the wild-type and ClpX I265V (or IGF::VGF) mutant were again minimized, only allowing the backbone of the IGF-loop to move in an unconstrained manner. Binding energies (ΔΔGbind) were calculated with the Rosetta energy function as ΔG complex – (ΔG ClpP, separated + ΔG ClpC/X,s eparated) using the Rosetta Interface Analyzer application, allowing side chains to re-pack in the separated interactions partners. To estimate the difference in binding in the *B. subtilis* ClpC-VGF and ClpC-IGF, the *B. subtilis* ClpP structure representing the open conformation (PDB ID: 3KTK) (*45*) was structurally aligned to 6SFX to have the *L. monocytogenes* ClpX IGF loops placed onto ClpP analogous to the *L. monocytogenes* ClpP-ClpX complex structure. Due to the symmetry mismatch between ClpP and ClpX, the six IGF loops were bound in distinct conformations, five of which are relatively similar (conformation 1), whereas one loop has a different conformation (conformation 2) (Fig S5E). In all conformations, changing Ile to Val diminishes the calculated binding energy (Fig S5E). Energy minimization and binding energy calculations were done as described above.

## Supporting information

supplemental Information for Alver et al

## Author contributions

RA, IH, NM, EC & KT were involved in conceptualization. RA, IH, FAC, KG & NM designed and performed the experiments. SR performed the *in silico* analysis for the binding energies. IH, RA, FAC & KT wrote the initial manuscript. All authors contributed to the manuscript writing, experimental design and analyzed the data. EC & KT supervised the study and were responsible for funding acquisition.

## Acknowledgements

We want to thank Jakob Fuhrmann (Scripps Research Institute) & Paul Thompson (University of Massachusetts Medical School) for samples of the anti-phospho-arginine antibody. David Dubnau (PHRI, Newark, NJ) & Axel Mogk (ZMBH, Uni Heidelberg) for strains and plasmids. Alex Elsholz (MPUSP, Berlin) for discussion. Christian Frese and Florian Kondrot (MPUSP, Berlin) for proteomic analysis.

