## supplemental Information for Alver et al for "The control of protein arginine phosphorylation facilitates proteostasis by an AAA+ chaperone protease system"

**Supplemental Figures S1-6 with Figure Legends (p2-13)**

**Supplemental Tables S1-S4 (p14-20)**

**References (p21)**

Figure S1

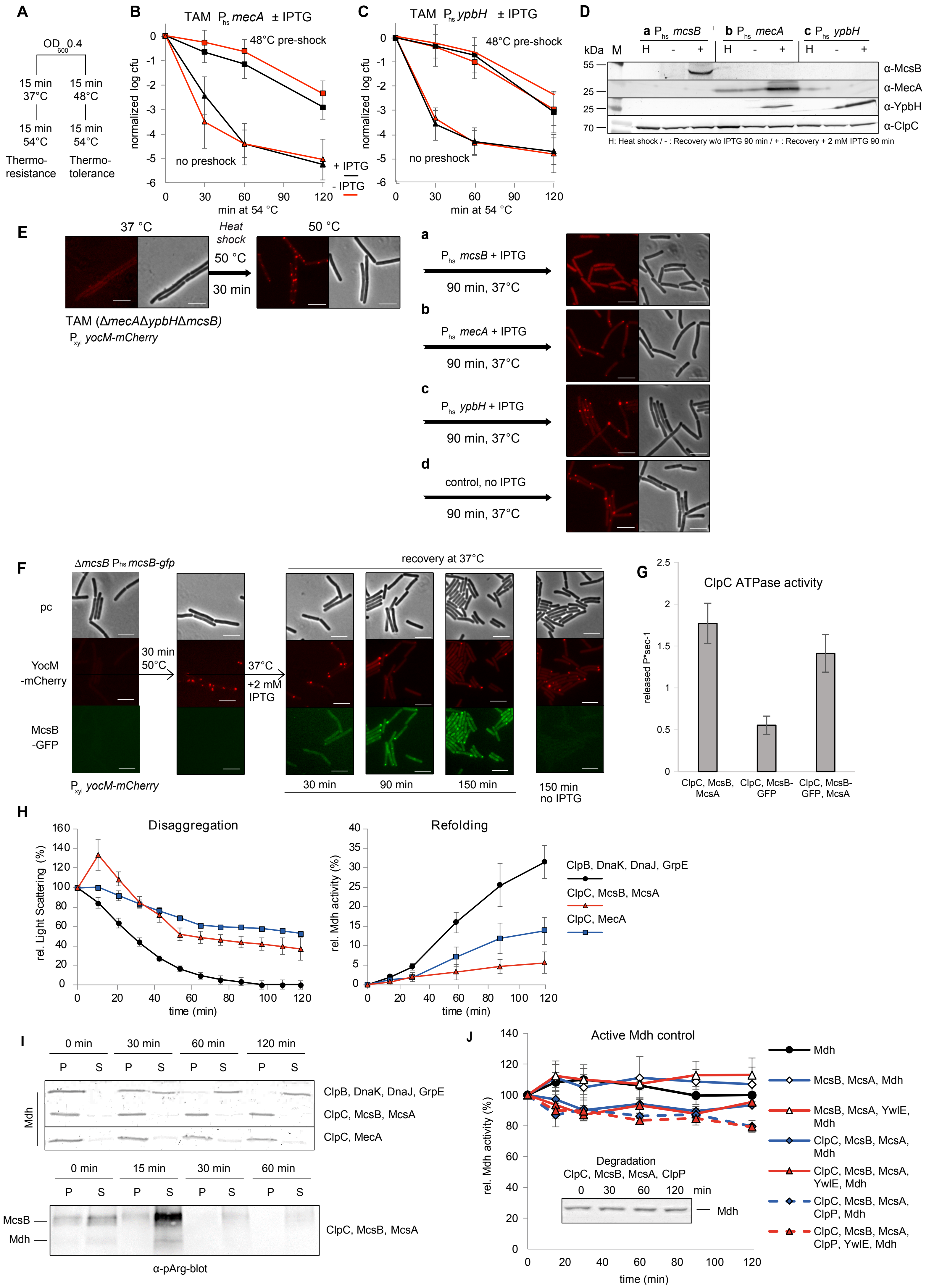
A
OD600 0.4
15 min 37°C
15 min 48°C
15 min 54°C
15 min 54°C
Thermo-resistance
Thermo-tolerance
B
TAM Phs mecA ± IPTG
48°C pre-shock
no preshock
normalized log cfu
min at 54 °C
+ IPTG
- IPTG
C
TAM Phs ypbH ± IPTG
48°C pre-shock
no preshock
normalized log cfu
min at 54 °C
D
a Phs mcsB
b Phs mecA
c Phs ypbH
kDa M
α-McsB
α-MecA
α-YpbH
α-ClpC
H: Heat shock / - : Recovery w/o IPTG 90 min / + : Recovery + 2 mM IPTG 90 min
E
37 °C
Heat shock
50 °C
30 min
50 °C
TAM (ΔmecAΔypbHΔmcsB)
Pxyl yocM-mCherry
a Phs mcsB + IPTG
90 min, 37°C
b Phs mecA + IPTG
90 min, 37°C
c Phs ypbH + IPTG
90 min, 37°C
d control, no IPTG
90 min, 37°C
F
ΔmcsB Phs mcsB-gfp
recovery at 37°C
pc
YocM -mCherry
McsB -GFP
30 min 50°C
37°C +2 mM IPTG
Pxyl yocM-mCherry
30 min
90 min
150 min
150 min no IPTG
G
ClpC ATPase activity
released P*sec-1
ClpC, McsB, McsA
ClpC, McsB-GFP
ClpC, McsB-GFP, McsA
H
Disaggregation
rel. Light Scattering (%)
time (min)
Refolding
rel. Mdh activity (%)
time (min)
ClpB, DnaK, DnaJ, GrpE
ClpC, McsB, McsA
ClpC, MecA
I
0 min
30 min
60 min
120 min
P
S
Mdh
ClpB, DnaK, DnaJ, GrpE
ClpC, McsB, McsA
ClpC, MecA
0 min
15 min
30 min
60 min
McsB
Mdh
ClpC, McsB, McsA
α-pArg-blot
J
Active Mdh control
rel. Mdh activity (%)
time (min)
Degradation
ClpC, McsB, McsA, ClpP
0 30 60 120 min
Mdh
McsB, McsA, Mdh
McsB, McsA, YwlE, Mdh
ClpC, McsB, McsA, Mdh
ClpC, McsB, McsA, YwlE, Mdh
ClpC, McsB, McsA, ClpP, Mdh
ClpC, McsB, McsA, ClpP, YwlE, Mdh

#### Figure S1 ClpC adaptor proteins

**A**) Scheme for the thermotolerance and thermoresistance experiment: At 37°C exponentially growing *B. subtilis* cells were either exposed to a mild heat shock for 15 min at 48°C before transfer to severe heat shock conditions at 54°C for 120 min, or they were directly exposed for 120 min to the severe 54°C growth conditions. Cell survival is measured by colony forming units (cfu) at the indicated time points. **B**) & **C**) Influence of MecA and YpbH on thermoresistance and thermotolerance. *B. subtilis* TAM $P_{hs}$ *mecA* strain (BIH740) and B) TAM + $P_{hs}$ *ypbH* strain (BIH741) were examined in a thermotolerance (squares: with pre-shock at 48 °C) and thermoresistance (triangles: no pre-shock) experiment, with (black) and without IPTG (red) to induce the expression of B) MecA or C) YpbH. $Log_{10}$-values of viable cell count were normalized. The data represents the mean of three biological replicates ± standard deviation (error bars). **D**) Adapter protein expression in complemented TAM strain. *B. subtilis* complemented TAM strains $P_{hs}$ *mcsB* (BIH739), $P_{hs}$ *mecA* (BIH740), $P_{hs}$ *ypbH* (BIH741) were grown in LB + 0,5 % xylose xylose (to induce the expression of the encoded $P_{xyl}$ *yocM*mCherry reporter), treated with a 50 °C heat shock and shifted to recover at 37 °C with 2 mM IPTG added for 90 min (see Fig 1D). Samples were taken for western blot analysis with Anti-McsB, Anti -MecA and Anti-YpbH antisera at indicated time points (after 30 min at 50°C "H" and after 90 min recovery at 37°C "-" without and "+" with IPTG). **E**) Subcellular protein aggregates detected by YocM-mCherry in the different *B. subtilis* complemented TAM strains (a: $P_{hs}$ *mcsB* (BIH739); b: $P_{hs}$ *mecA* (BIH740); c: $P_{hs}$ *ypbH* (BIH741); d: no IPTG control ($P_{hs}$ *ypbH* (BIH741)) were grown in LB with xylose, treated with a 50 °C heat shock for 30 min and shifted to recover for 90 min at 37 °C with IPTG. Samples were taken at indicated time points for inspection by fluorescence microscopy (scale bar 5 µm). **F**) McsB-GFP activity. *B. subtilis* strain Δ*mcsB* $P_{hs}$ *mcsB-gfp* $P_{xyl}$ *yocM-mCherry* (BIH671) was grown, treated with a 50 °C heat shock 30 min and subsequently shifted for 150 min to recover at 37 °C in the presence or absence of 2 mM IPTG. Samples were taken at indicated time points for fluorescence microscopy analysis with one representative image shown (scale bar 5 µm). **G**) McsB and McsB-GFP activated ClpC ATPase activity was determined by *in vitro* ATPase assay with three replicates to calculate standard deviations (error bars). **H**) Disaggregation and refolding experiments. Disaggregation of heat aggregated Mdh (1µM) after addition of ClpB/KJE (black circles), ClpC/McsB/McsA (red triangles) or ClpC/MecA (blue squares) was monitored *in vitro* by measuring light scattering at 30 °C for 120 min. The concurrent refolding was examined by measuring the Mdh enzymatic activity with samples from the disaggregation reaction taken at the indicated time points (right panel). 1,5 µM ClpB, 1 µM DnaK, 0,2 µM DnaJ, 0,1 µM GrpE, 1,5 µM ClpC, 1 µM MecA, 1 µM McsB, 1 µM McsA, 1 µM Mdh were used in all disaggregation and refolding experiments. **I**) Supernatant pellet experiments and protein arginine phosphorylation. Samples from the disaggregation and refolding experiment described in Fig 1F were taken at 0, 30, 60 and 120 min, centrifuged and the relative Mdh amount in the pellet (P) and supernatant (S) fractions was analyzed by Coomassie stained SDS-PAGE (upper panel). Samples from the disaggregation and refolding experiment with ClpC, McsB and McsA, described in Fig 1F, were taken at 0 min, 15 min, 30 min and analyzed by western blot experiments with anti-pArg antibody (Fuhrmann et al., 2015) (lower panel). **J**) Native Mdh activity in the presence of McsB kinase. Experiment measuring Mdh activity of native, not heat aggregated Mdh in presence of indicated protein combinations at 30 °C with the same buffer conditions and protein concentrations as in the presented disaggregation and refolding experiments (Fig 1F, 2A & 4). Inserted panel: Native Mdh stability in the presence of McsB kinase. ClpC/McsB/McsA/ClpP (1,5 /1/1/1 µM) were incubated with native Mdh (1µM) under the same buffer conditions and samples taken at the indicated timepoints by Coomassie stained SDS-PAGE analysis to assess the stability of native Mdh under these conditions in the presence of ClpP.

### Figure S2

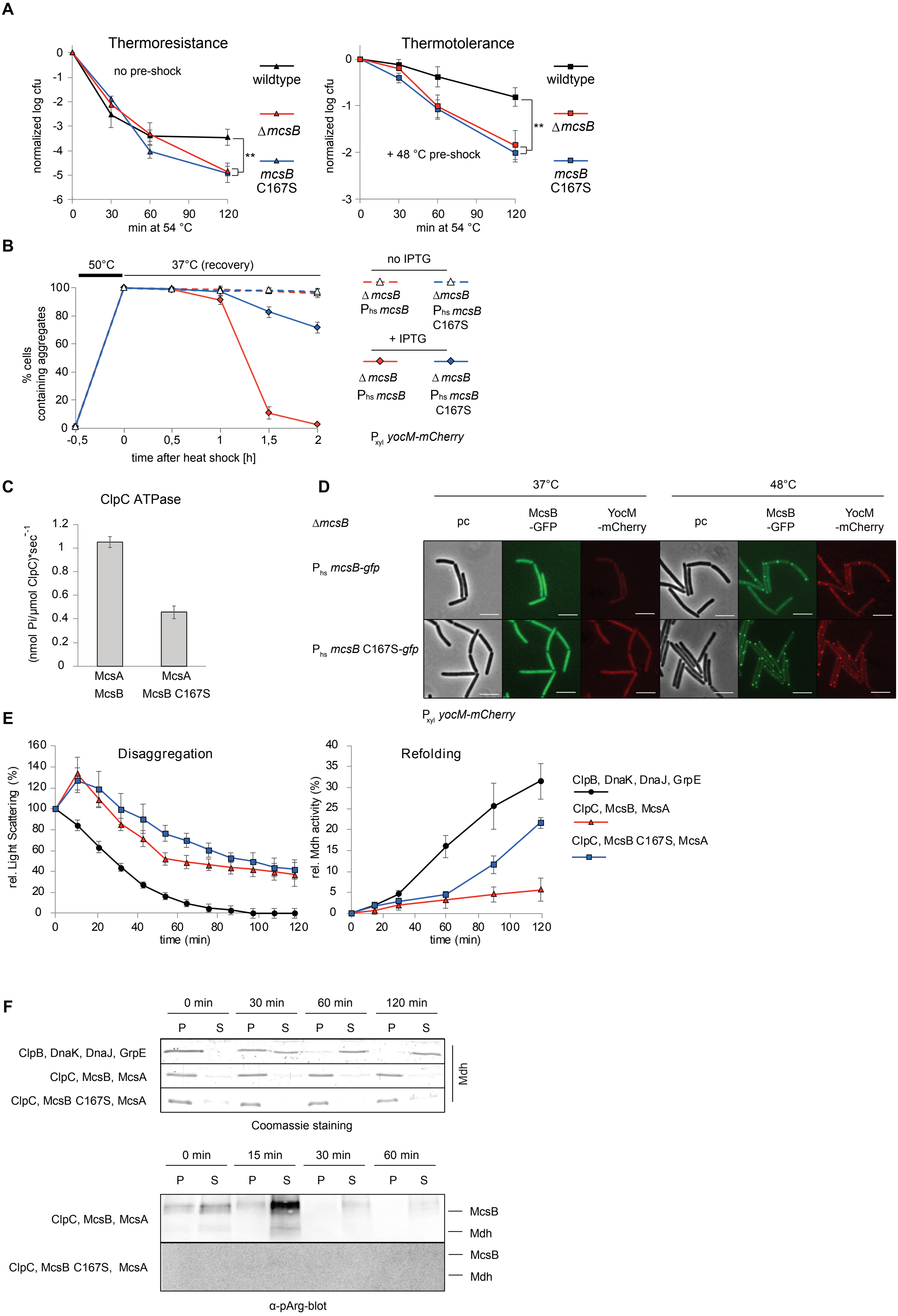
A
Thermoresistance
no pre-shock
normalized log cfu
min at 54 °C
wildtype
ΔmcsB
mcsB C167S
Thermotolerance
+ 48 °C pre-shock
B
50°C
37°C (recovery)
% cells containing aggregates
time after heat shock [h]
no IPTG
+ IPTG
ΔmcsB Phs mcsB
ΔmcsB Phs mcsB C167S
Pxyl yocM-mCherry
C
ClpC ATPase
(nmol Pi/µmol ClpC)*sec-1
McsA McsB
McsA McsB C167S
D
37°C
48°C
ΔmcsB
pc
McsB -GFP
YocM -mCherry
Phs mcsB-gfp
Phs mcsB C167S-gfp
Pxyl yocM-mCherry
E
Disaggregation
rel. Light Scattering (%)
time (min)
Refolding
rel. Mdh activity (%)
ClpB, DnaK, DnaJ, GrpE
ClpC, McsB, McsA
ClpC, McsB C167S, McsA
F
0 min
30 min
60 min
120 min
P
S
Mdh
Coomassie staining
15 min
McsB
α-pArg-blot

**Figure S2 Role of the McsB kinase activity in protein homeostasis**

**A**) Thermoresistance (left panel /triangles/ no pre-shock) and thermotolerance (right panel/squares: 15 min pre-shock at 48 °C) experiments with *B. subtilis* wildtype (black), Δ*mcsB* (BIH69, red) and *mcsB* C167S (BIH694, blue). $Log_{10}$-values of viable cell count (cfu) were normalized and three biological replicates were used to calculate standard deviations (error bars) with significant differences indicated (*: $p<0.05$, **: $p<0.01$, ***: $p<0.001$).

**B**) Decrease of subcellular protein aggregates during recovery after heat stress *in vivo*. *B. subtilis* Δ*mcsB* $P_{hs}$ *mcsB* $P_{xyl}$ *yocM-mCherry* (BIH414, red) and Δ*mcsB* $P_{hs}$ *mcsB* C167S $P_{xyl}$ *yocM-mCherry* (BIH439, blue) were grown in LB + 0,5 % xylose, treated with a 50 °C heat shock at $OD_{600}$ at 0.3 for 30 min and shifted to 37 °C in the absence or presence of 2 mM IPTG for 120 min. Samples were taken at indicated time points for analysis by fluorescence microscopy. To calculate the ratio of cells containing protein aggregates marked by YocM-mCherry, three biological replicates including three technical replicates with 50 – 100 cells were analyzed.

**C**) The *in vitro* ATPase activity of 1 µM ClpC with either 1 µM McsB or 1 µM McsB C167S in the presence of 1 µM McsA was determined. Three experimental replicates were carried out to calculate the standard deviations (error bars).

**D**) McsB and McsB-C167S co-localization with subcellular protein aggregates. *B. subtilis* strains Δ*mcsB* $P_{hs}$ *mcsB-gfp* $P_{xyl}$ *yocM-mCherry* (BIH671) and Δ*mcsB* $P_{hs}$ *mcsB* C167S-*gfp* $P_{xyl}$ *yocM-mCherry* (BIH672) were grown in LB in the presence of 0.5 % xylose and 1mM IPTG, allowing the expression of YocM-mCherry as marker for subcellular protein aggregates, in the presence of either McsB or McsB-C167S. These strains were grown to an $OD_{600}$ 0.4 at 37 °C and kept at 37°C or shifted to 48 °C for 15 min (left and right panel). Both were subsequently analyzed by fluorescence microscopy (representative images are shown) (scale bar 5 µm).

**E**) Disaggregation and refolding experiments. Disaggregation of heat aggregated Mdh after addition of ClpB/KJE (black circles), ClpC/McsB/McsA (red triangles) or ClpC/McsB C167S/McsA (blue squares) was monitored *in vitro* by measuring light scattering at 30 °C for 120 min (left panel). The concurrent Mdh refolding was examined by measuring the Mdh enzymatic activity at the indicated time points (right panel). McsB C167S and McsB were used at 1 µM.

**F**) Supernatant pellet experiments and protein arginine phosphorylation. Samples from the disaggregation and refolding experiment described in Fig 2C were taken at 0 min, 30 min, 60 min and 120 min, centrifuged and the relative Mdh amount in the pellet (P) and supernatant (S) fractions was analyzed by Coomassie stained SDS-PAGE (upper panel) (representative images are depicted). Samples from the disaggregation and refolding experiment with ClpC/ McsB/ McsA and ClpC/ McsB C167S/ McsA, described in Fig 2C, were taken at 0, 15, 30 & 60 min and analyzed by western blot experiments with anti-pArg antibody (Fuhrmann et al., 2015) (lower panel).

### Figure S3

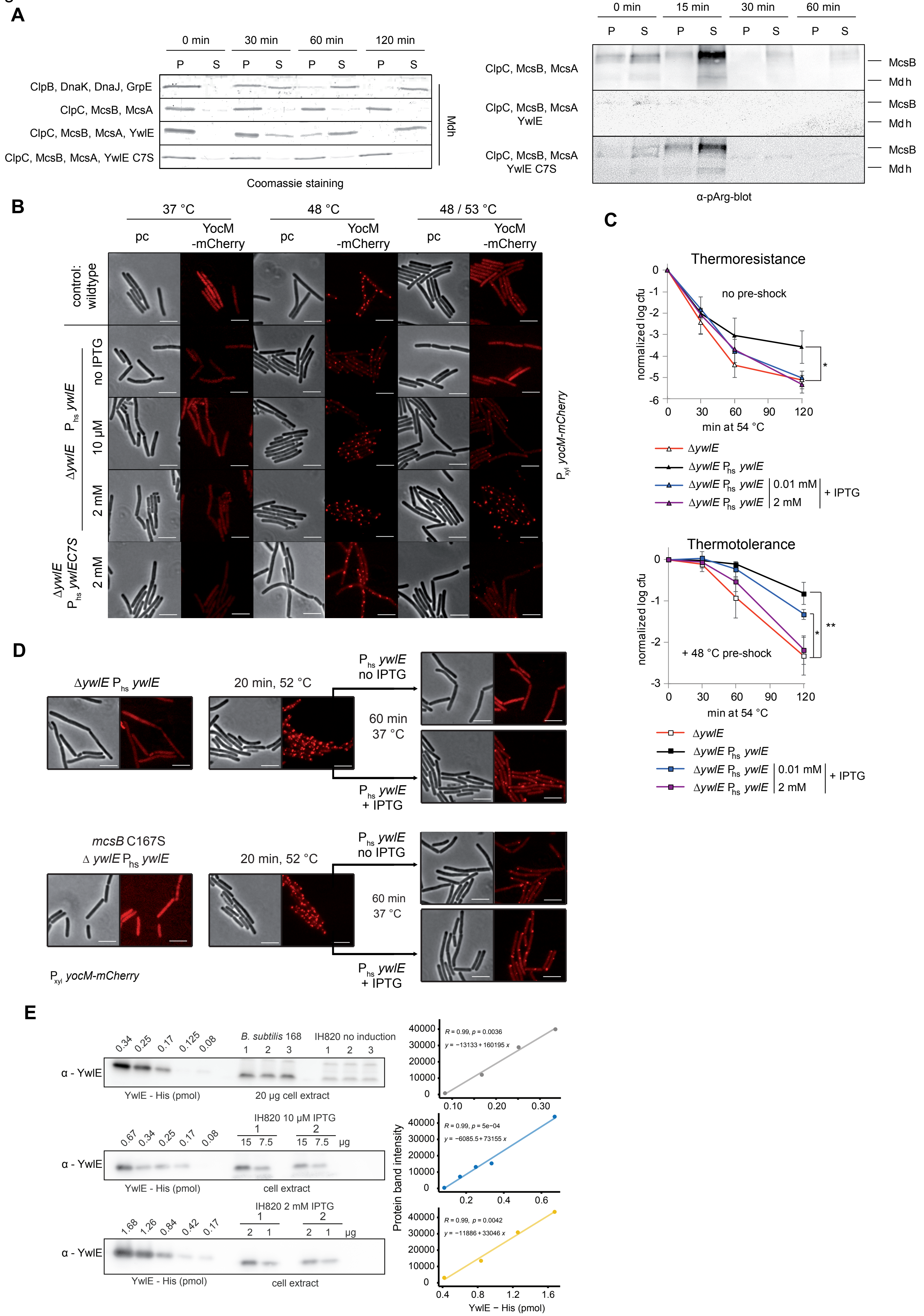
A
0 min
30 min
60 min
120 min
P S P S P S P S
ClpB, DnaK, DnaJ, GrpE
ClpC, McsB, McsA
ClpC, McsB, McsA, YwlE
ClpC, McsB, McsA, YwlE C7S
Mdh
Coomassie staining
0 min
15 min
30 min
60 min
ClpC, McsB, McsA
ClpC, McsB, McsA YwlE
ClpC, McsB, McsA YwlE C7S
McsB
Mdh
α-pArg-blot
B
37 °C
48 °C
48 / 53 °C
pc
YocM -mCherry
control: wildtype
no IPTG
10 µM
2 mM
ΔywlE $P_{hs}$ ywlE
ΔywlE $P_{hs}$ ywlEC7S
$P_{xyl}$ yocM-mCherry
C
Thermoresistance
no pre-shock
normalized log cfu
min at 54 °C
ΔywlE
ΔywlE $P_{hs}$ ywlE
ΔywlE $P_{hs}$ ywlE 0.01 mM
ΔywlE $P_{hs}$ ywlE 2 mM
+ IPTG
Thermotolerance
+ 48 °C pre-shock
D
ΔywlE $P_{hs}$ ywlE
20 min, 52 °C
$P_{hs}$ ywlE no IPTG
60 min 37 °C
$P_{hs}$ ywlE + IPTG
mcsB C167S
Δ ywlE $P_{hs}$ ywlE
$P_{xyl}$ yocM-mCherry
E
B. subtilis 168
IH820 no induction
α - YwlE
YwlE - His (pmol)
20 µg cell extract
IH820 10 µM IPTG
cell extract
IH820 2 mM IPTG
µg
R = 0.99, p = 0.0036
y = −13133 + 160195 x
R = 0.99, p = 5e−04
y = −6085.5 + 73155 x
R = 0.99, p = 0.0042
y = −11886 + 33046 x
Protein band intensity
YwlE − His (pmol)

#### Figure S3 YwlE and YwlEC7S

**A**) Supernatant pellet experiments and protein arginine phosphorylation. Samples from the disaggregation and refolding experiment described in Fig 3B were taken at 0, 30, 60 and 120 min, centrifuged and the Mdh amount in the pellet (P) and supernatant (S) fractions were analyzed by Coomassie stained SDS-PAGE (left panel) (representative images are depicted). Samples from the disaggregation and refolding experiment with the ClpC/McsB/McsA, ClpC/McsB/McsA/ YwlE and ClpC/McsB/McsA/YwlE C7S, as described in Fig 3B, were taken at 0, 15, 30 & 60 min and analyzed by western blot experiments with anti-pArg antibody (Fuhrmann et al., 2015) (right panel).

**B**) Visualization of subcellular protein aggregates *in vivo* under thermotolerance conditions in the presence of different amounts of YwlE. *B. subtilis* wildtype, Δ*ywlE* $P_{hs}$ *ywlE* (BIH828) and Δ*ywlE* $P_{hs}$ *ywlE* C7S (BIH829) cells also encoding $P_{xyl}$ *yocM-mCherry* were grown in LB + 0,5 % xylose and indicated concentrations of IPTG. Cells were exposed to the indicated temperatures. Samples were taken for analysis by fluorescence microscopy and representative pictures are shown (scale bar 5 µm).

**C**) Thermoresistance (upper panel no pre-shock) and thermotolerance (lower panel 15 min pre-shock at 48 °C) experiments with *B. subtilis* Δ*ywlE* (BIH309, red) and complemented Δ*ywlE* $P_{hs}$ *ywlE* (BIH828) with different YwlE expression levels due to the addition of different concentrations of IPTG (0/0.01/2 mM). $Log_{10}$-values of viable cell count were normalized, and three biological replicates were used to calculate standard deviations (error bars) with significant differences indicated (*: $p<0.05$, **: $p<0.01$, ***: $p<0.001$).

**D**) *B. subtilis* Δ*ywlE* $P_{hs}$ *ywlE* (BIH828) was grown in LB + 0.5 % xylose, treated with a 52 °C heat shock for 20 min at $OD_{600}$ at 0.3 and shifted to 37 °C in the absence or presence of 2 mM IPTG for 60 min and samples taken for subsequent analysis by fluorescence microscopy. *B. subtilis mcsB* C167S Δ*ywlE* $P_{hs}$ *ywlE* (BIH824) was grown in LB + 0.5 % xylose, treated with a 52 °C heat shock for 20 min at $OD_{600}$ of 0.3 and shifted to 37 °C for 60 min in the presence or absence of 2 mM IPTG with samples taken for fluorescence microscopy (lower panel). Representative fluorescence microscopy images for these experiments are depicted (scale bar 5 µm).

**E**) Exponentially growing cultures ($OD_{600}$ 0.35) of the Δ*ywlE* $P_{hs}$ *ywlE* (BIH820) strain were incubated with the indicated concentrations of IPTG in LB at 37 °C. Samples for SDS-PAGE and anti-YwlE western blot were taken after 30 min and the amount of cellular YwlE was determined by comparison with a calibration curve with MS-quantified His-tagged YwlE. Cell extracts protein concentrations were obtained by Bradford assay. Relative Western blot band intensity was acquired with ImageJ software and plotted against the respective sample protein amount. The linear regression used to determine each sample's YwlE amount is depicted next to the respective membrane. The number of cells for each sample was determined using a Petroff-Hauser chamber.

Figure S4

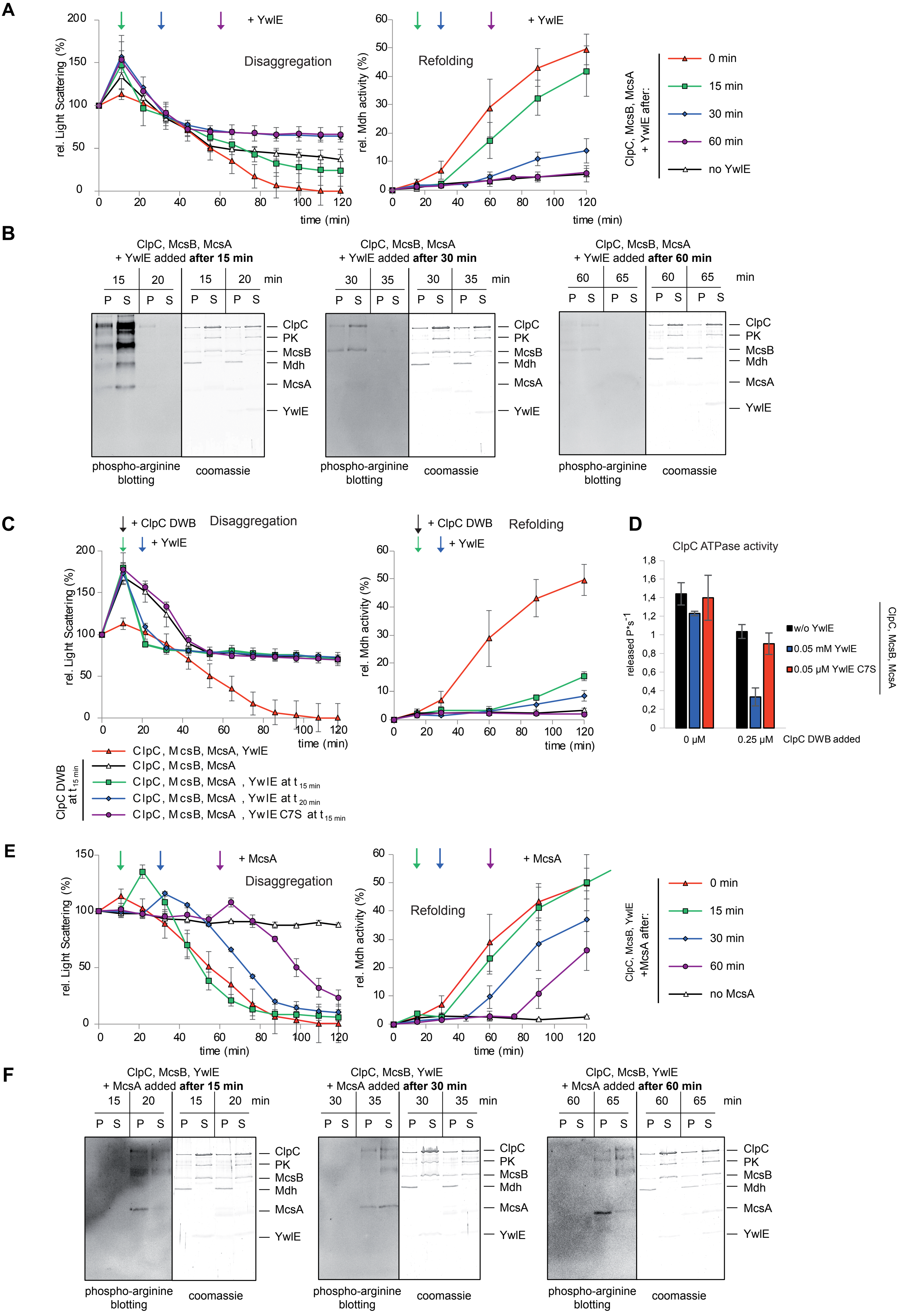

A
+ YwlE
Disaggregation
rel. Light Scattering (%)
time (min)
Refolding
rel. Mdh activity (%)
ClpC, McsB, McsA + YwlE after:
0 min
15 min
30 min
60 min
no YwlE
B
ClpC, McsB, McsA + YwlE added after 15 min
ClpC, McsB, McsA + YwlE added after 30 min
ClpC, McsB, McsA + YwlE added after 60 min
min
P S
ClpC
PK
McsB
Mdh
McsA
YwlE
phospho-arginine blotting
coomassie
C
+ ClpC DWB
+ YwlE
Disaggregation
Refolding
ClpC, McsB, McsA, YwlE
ClpC, McsB, McsA
ClpC, McsB, McsA , YwlE at t 15 min
ClpC, McsB, McsA , YwlE at t 20 min
ClpC, McsB, McsA , YwlE C7S at t 15 min
ClpC DWB at t 15 min
D
ClpC ATPase activity
released P*s-1
w/o YwlE
0.05 mM YwlE
0.05 µM YwlE C7S
ClpC, McsB, McsA
0 µM
0.25 µM
ClpC DWB added
E
+ McsA
Disaggregation
Refolding
ClpC, McsB, YwlE +McsA after:
0 min
15 min
30 min
60 min
no McsA
F
ClpC, McsB, YwlE + McsA added after 15 min
ClpC, McsB, YwlE + McsA added after 30 min
ClpC, McsB, YwlE + McsA added after 60 min
phospho-arginine blotting
coomassie

#### Figure S4 McsA & YwlE: kinase activation and de-phosphorylation

**A**) YwlE order of addition disaggregation and refolding experiments. Disaggregation (left panel) and refolding (right panel) of heat aggregated Mdh by ClpC/McsB in the presence of McsA, without (empty triangles, black) and with the addition of YwlE at different timepoints (0 min (triangle, red), 15 min (square, green), 30 min (diamond, blue), 60 min (circle, purple)). The refolding of Mdh was examined by measuring the Mdh enzymatic activity with samples taken at indicated different time points (right panel). At least three experiments were implemented, and the observed standard deviation indicated as error bars.

**B**) Samples from the disaggregation and refolding experiment with ClpC/McsB/McsA, with YwlE added at 0, 15, 30 & 60 min described in Fig S4A were taken just before and 5 min after addition of YwlE and analyzed by western blot experiments with anti-pArg antibody (Fuhrmann et al., 2015) (left panel). The corresponding Coomassie-stained SDS-PAGE of pellet (P) and supernatant (S) fractions is depicted in the right panel. The respective proteins are indicated on the right (PK: pyruvate kinase).

**C**) ClpC-DWB & YwlE. Disaggregation (left panel) and refolding (right panel) of heat aggregated Mdh by ClpC, McsB & McsA adding 0.25 μM ClpC DWB after 15 min (empty triangles, black) and YwlE after 15 min or 20 min (squares, green & diamonds, black). In another experiment ClpC-DWB and YwlE C7S were added after 15 min to ClpC, McsB & McsA. The disaggregation and refolding with ClpC, McsB, McsA & YwlE (triangles, red) is shown as the positive control. The refolding of Mdh was examined by measuring the enzymatic activity at indicated time points (right panel).

**D**) Inhibition of *in vitro* ClpC ATPase activity by addition of ClpC-DWB (0.25μM) to ClpC/McsB/McsA (1/1/1 μM), in the presence or absence of either YwlE or YwlE C7S (0.05 μM).

**E**) McsA order of addition disaggregation and refolding experiments. Disaggregation (left panel) and refolding (right panel) of heat aggregated Mdh by ClpC/McsB in the presence of YwlE, without (triangles black) and with the addition of McsA at different timepoints (0 min (triangle, red), 15 min (square green), 30 min (diamond, blue), 60 min (circle purple)). The refolding of Mdh was examined by measuring the Mdh enzymatic activity with samples taken at indicated time points (right panel).

**F**) Samples from the order of addition disaggregation and refolding experiment with ClpC/McsB/YwlE, and McsA added at 0, 15, 30 & 60 min described in Fig 3B were taken just before and 5 min after addition of McsA and analyzed by western blot experiments with anti-pArg antibody (Fuhrmann et al., 2015) (left panel) The corresponding Coomassie-stained SDS-PAGE of the analyzed pellet (P) and supernatant (S) fractions is depicted in the right panel.

### Suppl Figure S5

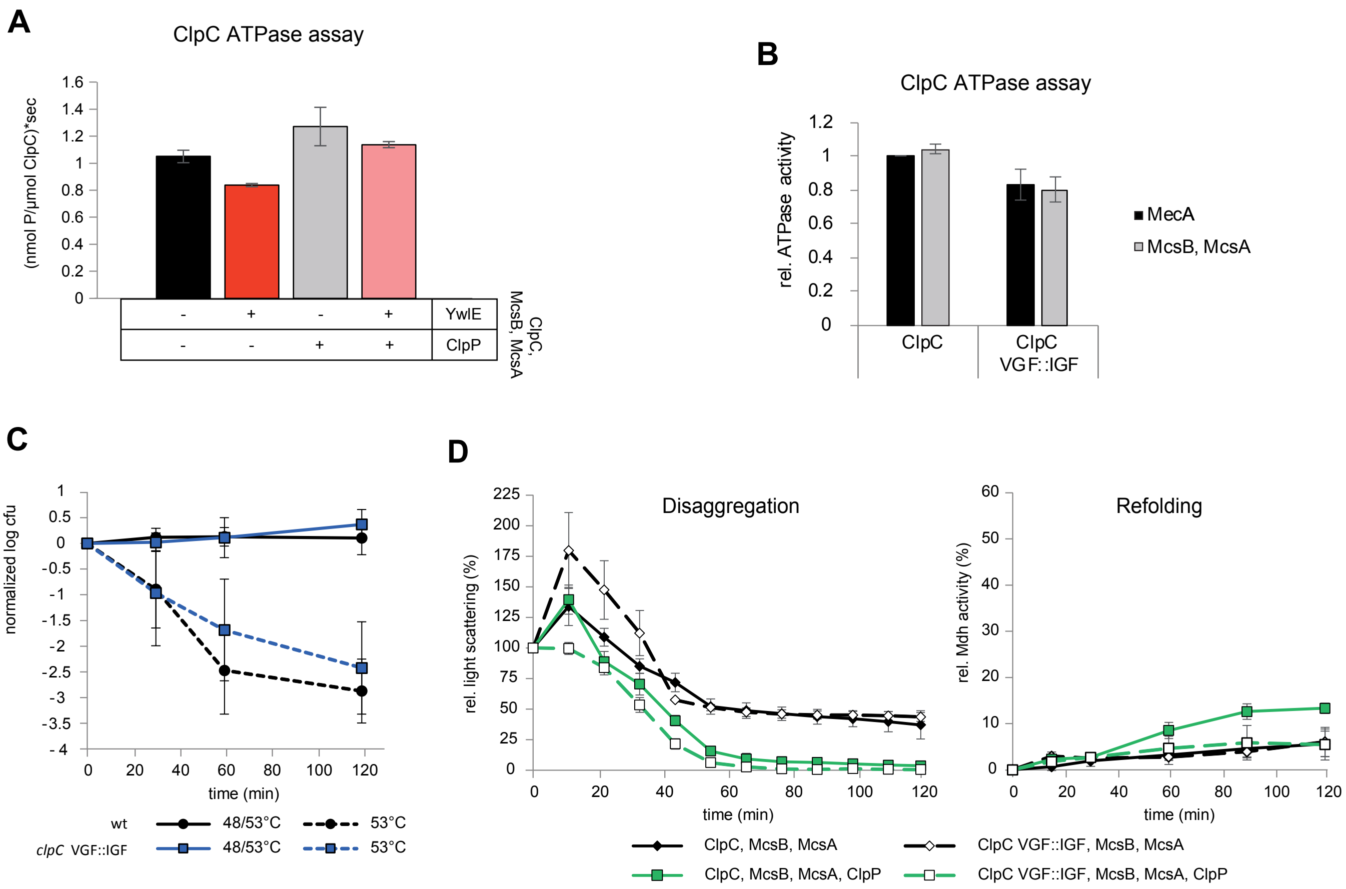


**E**

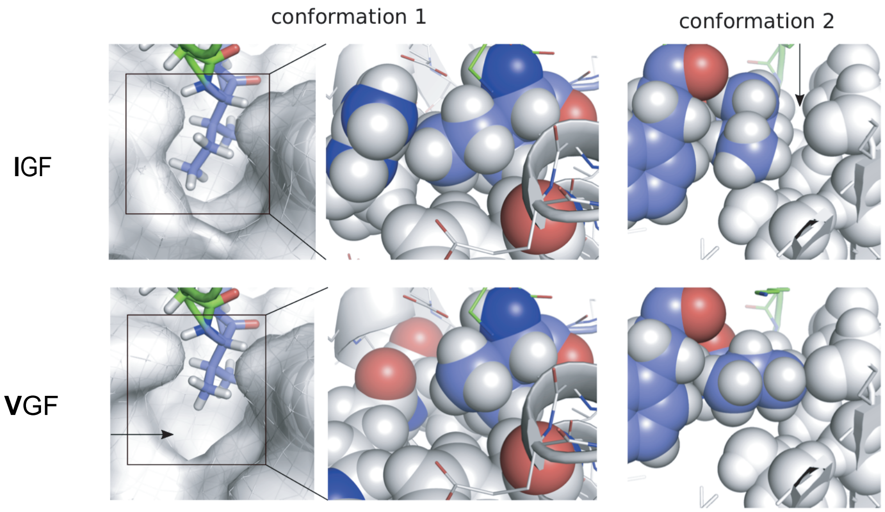


*Bacillus subtilis*

| Config. | IGF | VGF | $\Delta\Delta G_{bind}$ |
|---|---|---|---|
| 1a | -17.1 | -18.6 | 1.5 |
| 1b | -20.8 | -18.4 | -2.4 |
| 1c | -19.3 | -18.2 | -1.1 |
| 1d | -32.0 | -25.9 | -6.1 |
| 1e | -30.8 | -35.6 | -3.2 |
| 2 | -18.0 | -19.1 | 1.1 |
| sum | -146.0 | -135.8 | -10.2 |

*Listeria mon.*

| Config. | IGF | VGF | $\Delta\Delta G_{bind}$ |
|---|---|---|---|
| 1a | -50.5 | -45.7 | -4.7 |
| 1b | -46.2 | -38.7 | -7.5 |
| 1c | -38.0 | -37.9 | -0.1 |
| 1d | -42.3 | -34.4 | -8.0 |
| 1e | -44.9 | -41.0 | -3.0 |
| 2 | -36.2 | -31.8 | -4.4 |
| sum | -258.1 | -230.4 | -27.7 |

#### Figure S5 P-loop mutation

**A**) ClpC ATPase assay in the presence of McsB & McsA, ±YwlE and ClpP (all at 1µM). Error bars represent the standard deviation of three replicates.

**B**) *In vitro* ATPase activity of 1 µM ClpC and 1 µM ClpC VGF::IGF with either 1 µM MecA or 1 µM McsB & McsA. Error bars represent the standard deviation of three replicates.

**C**) Thermoresistance and thermotolerance *B. subtilis* wildtype (filled circles black) and *clpC* VGF::IGF (BRK166, filled squares, blue) were investigated in a thermoresistance (dotted lines, no pre-shock) and thermotolerance experiment (Solid line: + 15 min pre-shock at 48 °C). $Log_{10}$-values of viable cell count were normalized, and three biological replicates were used to calculate standard deviations (error bars).

**D**) Disaggregation of heat aggregated Mdh (1µM) after addition of ClpC/ McsB/ McsA (filled diamonds, black) ClpC/ McsB/ McsA/ ClpP (filled squares, green), ClpC VGF::IGF/ McsB/ McsA (empty diamonds, black) and ClpC VGF::IGF/ McsB/ McsA/ ClpP (empty squares, black, green line) was monitored *in vitro* by measuring the decrease of light scattering at 30 °C for 120 min (left panel). The concurrent refolding was examined by measuring the Mdh enzymatic activity with samples from the disaggregation reaction taken at the indicated time points (right panel).

**E**) Rosetta modeling software was used to perform energy minimization of each of the six ClpP-bound IGF loops and compared the calculated binding energies between the wild type and an Ile-to-Val in silico exchange. Left: Model of P-loop bound to *B. subtilis* ClpP. Comparison of IGF/VGF loops (light blue) when bound to *B. subtilis* ClpP (white) in the two major conformations as found in the *L. monocytogenes* ClpX-ClpP complex structure. Conformation 1 is shown in stick and sphere representation; conformation 2 is shown as spheres. Arrows indicate missing contacts. Right: Calculated Rosetta binding energies for all six conformations. Top: IGF-loop (backbone from *L. monocytogenes* ClpX) docked into *B. subtilis* ClpP Bottom: IGF-loop from *L. monocytogenes* bound to *L. monocytogenes* ClpP. $\Delta\Delta G_{bind}$ denotes the difference in binding energy between IGF and VGF loops.

### Figure S6

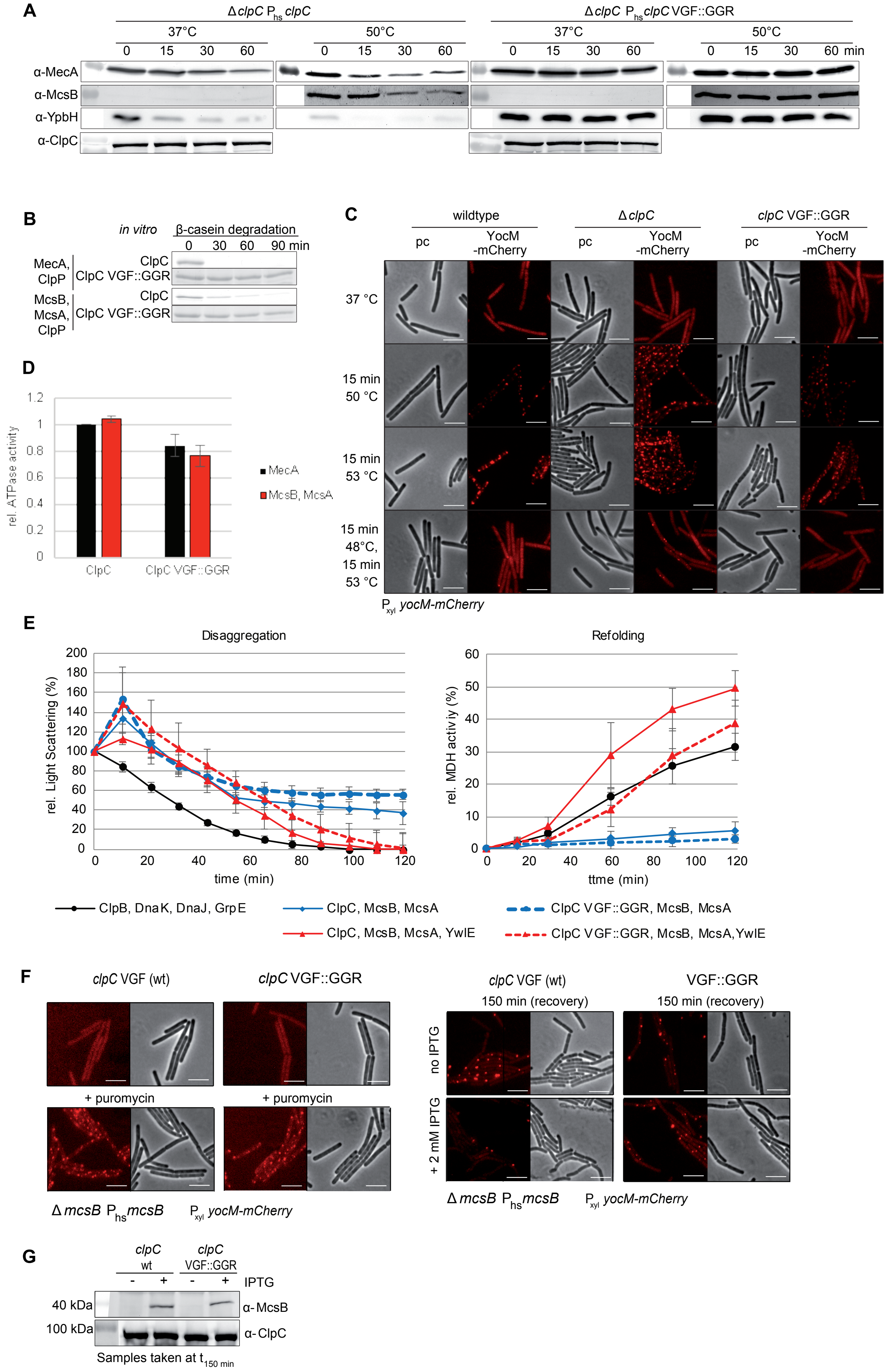
A
ΔclpC Phs clpC
ΔclpC Phs clpC VGF::GGR
37°C
50°C
0 15 30 60
0 15 30 60 min
α-MecA
α-McsB
α-YpbH
α-ClpC
B
in vitro
β-casein degradation
0 30 60 90 min
MecA, ClpP
ClpC
ClpC VGF::GGR
McsB, McsA, ClpP
C
wildtype
ΔclpC
clpC VGF::GGR
pc
YocM -mCherry
37 °C
15 min 50 °C
15 min 53 °C
15 min 48°C, 15 min 53 °C
Pxyl yocM-mCherry
D
rel. ATPase activity
ClpC
ClpC VGF::GGR
MecA
McsB, McsA
E
Disaggregation
Refolding
rel. Light Scattering (%)
rel. MDH activity (%)
time (min)
ClpB, DnaK, DnaJ, GrpE
ClpC, McsB, McsA
ClpC VGF::GGR, McsB, McsA
ClpC, McsB, McsA, YwlE
ClpC VGF::GGR, McsB, McsA, YwlE
F
clpC VGF (wt)
clpC VGF::GGR
+ puromycin
ΔmcsB Phs mcsB
Pxyl yocM-mCherry
VGF::GGR
150 min (recovery)
no IPTG
+ 2 mM IPTG
G
clpC wt
clpC VGF::GGR
IPTG
40 kDa
100 kDa
α-McsB
α-ClpC
Samples taken at t150 min

**Figure S6 ClpC VGF::GGR loop mutant**

**A**) *in vivo* stability of adaptor proteins *B. subtilis* strain Δ*clpC* $P_{hs}$ *clpC* (BIH152) and Δ*clpC* $P_{hs}$ *clpC* VGF::GGR (BIH504) were grown at 37 °C and 50 °C in LB medium + 1 mM IPTG until $OD_{600}$ 0.4 and treated with 25 µg/mL chloramphenicol (f.c.) to arrest new protein synthesis. Samples were taken for western blot with indicated anti-MecA, anti-McsB, anti-YpbH & anti-ClpC antibodies at indicated time points.

**B**) *In vitro* degradation of β-casein at 37 °C by either 1 µM ClpC & 1 µM ClpP or 1 µM ClpC VGF::GGR & 1 µM ClpP in combination with 1 µM MecA or 1 µM McsB & 1 µM McsA. Samples for Coomassie stained SDS-PAGE analysis of β-casein were taken after 0 min, 30 min, 60 min and 120 min.

**C**) *B. subtilis* wildtype, Δ*clpC* (BIH865) and *clpC* VGF::GGR (BIH792) were grown in LB medium and treated at $OD_{600}$ 0.4 according to indicated thermoresistance and thermotolerance conditions. Samples were taken for analysis of the YocM-mCherry marked subcellular protein aggregates by fluorescence microscopy. Representative images are depicted (scale bar 5 µm).

**D**) *In vitro* ATPase activity of ClpC and ClpC VGF::GGR with either MecA or McsB/McsA (all at 1µM). Error bars represent the standard deviation of three replicates.

**E**) Disaggregation of heat aggregated 1µM Mdh after addition of ClpBKJE (filled circles, black) ClpC/McsB/McsA (filled diamonds, blue), ClpC/McsB/McsA/YwlE (filled triangles, red), ClpC VGF::GGR/McsB/McsA (filled circles, blue) and ClpC VGF::GGR/McsB/McsA/YwlE (filled triangles, red, dotted line) was monitored *in vitro* by measuring the decrease of light scattering at 30 °C for 120 min (left panel). The concurrent refolding was examined by measuring the Mdh enzymatic activity at the indicated time points (right panel).

**F**) *B. subtilis* clpC wt (Δ*mcsB* $P_{hs}$ *mcsB* $P_{xyl}$ *yocM-mCherry* (BIH414)) and *clpC* VGF::GGR Δ*mcsB* $P_{hs}$ *mcsB* $P_{xyl}$ *yocM-mCherry* (BIH485)) were grown in LB in the presence of 0.5 % xylose and treated with 25 µg/mL puromycin. After 15 min, puromycin was removed by centrifugation and resuspension of the cells in fresh media with xylose in the presence or absence of 2 mM IPTG. After incubation at 37 °C for 150 min, samples were taken for analysis by fluorescence microscopy.

**G**) Western blot analysis of *B. subtilis* Δ*mcsB* $P_{hs}$ *mcsB* $P_{xyl}$ *yocM-mCherry* (BIH414) and *clpC* VGF::GGR Δ*mcsB* $P_{hs}$ *mcsB* $P_{xyl}$ *yocM-mCherry* (BIH485) strains with anti-McsB or anti-ClpC antibodies after 150 min recovery from puromycin treatment in the presence or absence of IPTG (see Fig 5D, FigS6F)

#### Supplementary tables

**Table S1:** List of strains

| Strains | Relevant genotype/properties | Source/construction |
|---|---|---|
| ***E. coli* DH5α** | $F^-$ Φ80 lacZΔM15 Δ(lacZYA-*arg*F) U169 *rec*A1 *end*A1 *hsd*R17($r_k^-$, $m_k^+$) *pho*A *sup*E44 *thi*-1 *gyr*A96 *rel*A1 $λ^-$ | Invitrogen/ThermoFischerScientific |
| ***E. coli* BL21(DE3)** | $F^–$ *omp*T *hsd*$S_B$ ($r_B^–$, $m_B^–$) *dcm gal* λ(DE3) pLysS $Cm^r$ | Invitrogen/ThermoFischerScientific |
| ***B. subtilis* 168** | *trpC2* | (Anagnostopoulos and Spizizen, 1961) |
| **QBP418** | PY79 Δ*clpC::tet* | (Pan et al., 2001) |
| **BIH19** | *trpC2* Δ*clpC::tet* | QBP418 → *B. subtilis* 168 |
| **BEK89** | *trpC2 lys-3* Δ*mcsB::kan* | (Krüger et al., 2001) |
| **BIH69** | *trpC2* Δ*mcsB::kan* | BEK89 → *B. subtilis* 168 |
| **BIH73** | *trpC2 amyE::*$P_{xyl}$ *yocM-mCherry spec* | (Hantke et al., 2019) |
| **BIH140** | *trpC2 clpC* E280A E618A (DWB) | (Kirstein et al., 2006) |
| **BIH151** | *trpC2 amyE::*$P_{hs}$ *clpC spec* | Plasmid 6 → BIH1 |
| **BIH152** | *trpC2 amyE::*$P_{hs}$ *clpC spec* Δ*clpC::tet* | BIH19 → BIH151 |
| **BIH217** | *trpC2 clpC* VGF::GGR | (Moliere, 2012) |
| **BIH309** | *trpC2* Δ*ywlE::kan* | (Elsholz et al., 2010) |
| **BIH369** | *trpC2 lacA::*$P_{xyl}$ *yocM-mCherry ery* | (Hantke et al., 2019) |
| **BIH399** | *trpC2 mcsB::kan lacA::*$P_{xyl}$ *yocM-mcherry ery* | BIH69 → BIH369 |
| **BIH407** | *trpC2 amyE::*$P_{hs}$ *mcsB spec clpC* VGF::GGR | Plasmid 8 → BIH217 |
| **BIH411** | *trpC2 amyE::*$P_{hs}$ *mcsB spec clpC* E280A E618A (DWB) | Plasmid 8 → BIH140 |
| **BIH414** | *trpC2 amyE::*$P_{hs}$ *mcsB spec mcsB::kan lacA::*$P_{xyl}$ *yocM-mcherry ery* | Plasmid 8 → BIH399 |
| **BIH423** | *trpC2 amyE::*$P_{hs}$ *mcsB spec mcsB::kan clpC* VGF::GGR | BIH69 → BIH407 |
| **BIH427** | *trpC2 amyE::*$P_{hs}$ *mcsB spec mcsB::kan clpC* E280A E618A (DWB) | BIH69 → BIH411 |

| | | |
|---|---|---|
| **BIH432** | *trpC2 lacA::$P_{xyl}$ yocM-mCherry ery amyE::$P_{hs}$ clpC spec ΔclpC::tet* | (Hantke et al., 2019) |
| **BIH434** | *trpC2 amyE::$P_{hs}$ clpC* VGF::GGR *spec lacA::$P_{xyl}$ yocM-mCherry ery* | Plasmid 7 → BIH369 |
| **BIH439** | *trpC2 amyE::$P_{hs}$ mcsB* C167S *spec mcsB::kan lacA::$P_{xyl}$ yocM-mcherry ery* | Plasmid 12 → BIH399 |
| **BIH485** | *trpC2 amyE::$P_{hs}$ mcsB spec mcsB::kan lacA::$P_{xyl}$ yocM-mcherry ery clpC* VGF::GGR | BIH369 → BIH423 |
| **BIH488** | *trpC2 amyE::$P_{hs}$ mcsB spec mcsB::kan lacA::$P_{xyl}$ yocM-mcherry ery clpC* E280A E618A (DWB) | BIH369 → BIH427 |
| **BIH504** | *trpC2 amyE::$P_{hs}$ clpC* VGF::GGR *spec ΔclpC::tet lacA::$P_{xyl}$ yocM-mCherry ery* | BIH19 → BIH434 |
| **BIH662** | *trpC2 amyE::$P_{hs}$ mcsB-gfp* A206K *spec* | Plasmid 11 → BIH1 |
| **BIH663** | *trpC2 amyE::$P_{hs}$ mcsB* C167S *-gfp* A206K *spec* | Plasmid 13 → BIH1 |
| **BIH671** | *trpC2 lacA::$P_{xyl}$ yocM-mCherry ery amyE::$P_{hs}$ mcsB-gfp* A206K *spec* | BIH369 → BIH662 |
| **BIH672** | *trpC2 lacA::$P_{xyl}$ yocM-mCherry ery amyE::$P_{hs}$ mcsB* C167S *-gfp* A206K *spec* | BIH369 → BIH663 |
| **BIH694** | *trpC2 mcsB::mcsB* C167S | Plasmid 16 → BIH1 |
| **BIH695** | *trpC2 mcsB::mcsB* C167S *ywlE::kan* | *ywlE::kan* (PCR) → BIH694 |
| **BIH721** | *trpC2 lacA::$P_{xyl}$ yocM-mCherry ery ΔypbH::cat* | *ypbH::cat* (PCR) → BIH369 |
| **BIH735** | *trpC2 lacA::$P_{xyl}$ yocM-mCherry ery ΔypbH::cat ΔmcsB::kan* | BIH69 → BIH721 |
| **BIH737** | *trpC2 lacA::$P_{xyl}$ yocM-mCherry ery ΔypbH::cat ΔmcsB::kan ΔmecA::tet* | BIH20 → BIH735 |
| **BIH739** | *trpC2 lacA::$P_{xyl}$ yocM-mCherry ery ΔypbH::cat ΔmcsB::kan ΔmecA::tet amyE::$P_{hs}$ mcsB spec* | Plasmid 8 → BIH737 |
| **BIH740** | *trpC2 lacA::$P_{xyl}$ yocM-mCherry ery ΔypbH::cat ΔmcsB::kan ΔmecA::tet amyE::$P_{hs}$ mecA spec* | Plasmid 9 → BIH737 |
| **BIH741** | *trpC2 lacA::$P_{xyl}$ yocM-mCherry ery ΔypbH::cat ΔmcsB::kan ΔmecA::tet amyE::$P_{hs}$ ypbH spec* | Plasmid 10 → BIH737 |
| **BIH792** | *trpC2 lacA::$P_{xyl}$ yocM-mCherry ery clpC::clpC* VGF::GGR (671-673) | BIH369 → BIH217 |

| | | |
|---|---|---|
| **BIH805** | *trpC2 mcsB::mcsB* C167S *ywlE::kan lacA::$P_{xyl}$ yocM-mcherry ery* | BIH695 → BIH369 |
| **BIH816** | *trpC2 amyE::$P_{hs}$ ywlE spec* | Plasmid 14 → BIH309 |
| **BIH817** | *trpC2 amyE::$P_{hs}$ ywlE* C7S *spec* | Plasmid 15 → BIH309 |
| **BIH819** | *trpC2 lacA::$P_{xyl}$ yocM-mCherry ery ΔywlE::kan* | BIH369 → BIH309 |
| **BIH820** | *trpC2 amyE::$P_{hs}$ ywlE spec ΔywlE::kan* | BIH309 → BIH816 |
| **BIH821** | *trpC2 amyE::$P_{hs}$ ywlE* C7S *spec ΔywlE::kan* | BIH309 → BIH817 |
| **BIH824** | *trpC2 mcsB::mcsBC167S ywlE::kan lacA::$P_{xyl}$ yocM-mcherry ery amyE::$P_{hs}$ ywlE spec* | BIH805 → BIH816 |
| **BIH828** | *trpC2 lacA::$P_{xyl}$ yocM-mCherry ery amyE::$P_{hs}$ ywlE spec ΔywlE::kan* | BIH369 → BIH820 |
| **BIH829** | *trpC2 lacA::$P_{xyl}$ yocM-mCherry ery amyE::$P_{hs}$ ywlE* C7S *spec ΔywlE::kan* | BIH369 → BIH821 |
| **BIH860** | *trpC2 clpC VGF::IGF mcsB::kan amyE::Plac mcsB spec lacA::Pxyl yocM-mCherry ery* | |
| **BIH865** | *trpC2 lacA::$P_{xyl}$ yocM-mCherry ery ΔclpC::tet* | BIH369 → BIH19 |
| **BRK166** | *trpC2 clpC* VGF::IGF | Plasmid 28 → *B. subtilis* 168 |
| **ERK55** | *pET28a clpP* | Plasmid 23 → BL21 (D3) |
| **ERK68** | *pET28a clpC* | Plasmid 17 → BL21 (D3) |
| **ERK75** | *pCA528 mcsB* | Plasmid 19 → BL21 (D3) |
| **ERK95** | *pQE32 mcsA* | Plasmid 18 → BL21 (D3) |
| **ERK106** | *pET28a mecA* | Plasmid 21 → BL21 (D3) |
| **ERK110** | *pET28a clpC* VGF::GGR | Plasmid 24 → BL21 (D3) |
| **ERK116** | *pET28a clpC* VGF::IGF | Plasmid 25 → BL21 (D3) |
| **ERK164** | *pET28a ywlE* | Plasmid 22 → BL21 (D3) |
| **ERK195** | *pCA528 mcsB* C167S | Plasmid 20 → BL21 (D3) |
| **ERK198** | *pET28a clpC* E280A E618A | Plasmid 26 → BL21 (D3) |
| **ERK315** | *pDS56 clpB- placIq* | (Mogk) |
| **ERK316** | *pSUMO grpE* | (Mogk) → BL21 (D3) |
| **ERK318** | *pSUMO dnaK* | (Mogk) → BL21 (D3) |

| | | |
|---|---|---|
| **ERK319** | *pSUMO dnaJ* | (Mogk) → BL21 (D3) |
| **ERK441** | *pET28a ywlE C7S* | Plasmid 27→ BL21 (D3) |
| **EIH798** | *pQE60 ywlE* C7S | Plasmid 27 → BL21 (D3) |

**Table S2: List of plasmids**

| Nr. | Plasmid | Construction | Reference |
|---|---|---|---|
| **1** | pSG1154-*gfp*A206K | *amyE*-integrating plasmid, xylose inducible, *cat*$^R$ | (Lewis and Marston, 1999) |
| **2** | pSG1154 *yocM-mCherry* | *yocM-mCherry* in pSG1154 | (Hantke et al., 2019) |
| **3** | pBS2E | *lacA*-integrating plasmid, *ery*$^R$ | (Radeck et al., 2013) |
| **4** | pBS2E like pDR111 | $P_{hs}$ *gene of interest* + *lacI* in pBS2E backbone, *ery*$^R$ | (Hantke et al., 2019) |
| **5** | pDR111 | *amyE*-integrating plasmid, IPTG inducible, *spec*$^R$ | Provided by David Dubnau |
| **6** | pDR111 *clpC* | *clpC* in pDR111 | (Hantke et al., 2019) |
| **7** | pDR111 *clpC* VGF::GGR | *clpC* in pDR111 | This work, template from (Moliere, 2012) |
| **8** | pDR111 *mcsB* | Expression of *mcsB*, $P_{hs}$ promoter, *spec*$^R$ | This work |
| **9** | pDR111 *mecA* | Expression of *mecA*, $P_{hs}$ promoter, *spec*$^R$ | This work |
| **10** | pDR111 *ypbH* | Expression of *ypbH*, $P_{hs}$ promoter, *spec*$^R$ | This work |
| **11** | pDR111 *mcsB-gfp* | Expression of *mcsB-gfp*A206K, $P_{hs}$ promoter, *spec*$^R$ | This work |
| **12** | pDR111 *mcsB* C167S | Expression of *mcsB* C167S, $P_{hs}$ promoter, *spec*$^R$ | This work |
| **13** | pDR111 *mcsB* C167S-*gfp* | Expression of *mcsB* C167S-*gfp* A206K, $P_{hs}$ promoter, *spec*$^R$ | This work |

| | | | |
|---|---|---|---|
| **14** | pDR111 *ywlE* | Expression of *ywlE*, $P_{hs}$ promoter, *spec*$^R$ | This work |
| **15** | pDR111 *ywlE* C7S | Expression of *ywlE* C7S, $P_{hs}$ promoter, *spec*$^R$ | This work |
| **16** | pMAD *mcsB* C167S | For *B. subtilis* markerless point mutation *mcsB* C167S *in cis*, *ery*$^R$ | This work |
| **17** | pET28a *clpC* | Expression of *B. subtilis* ClpC with $His_6$-tag | This work |
| **18** | pQE32 *mcsA* | Expression of *B. subtilis* McsA with His6-tag | (Kirstein, 2005) |
| **19** | pCA528 *mcsB* | Expression of *B. subtilis* McsB with SUMO-$His_6$-tag | (Kirstein, 2005) |
| **20** | pCA528 *mcsB* C167S | Expression of *B. subtilis* McsB C167S with SUMO-$His_6$-tag | This work |
| **21** | pET28a *mecA* | Expression of B. subtilis MecA with $His_6$-tag | This work |
| **22** | pET28a *ywlE* | Expression of B. subtilis YwlE with $His_6$-tag | This work |
| **23** | pET28a *clpP* | Expression of B. subtilis ClpP with $His_6$-tag | This work |
| **24** | pET28a c*lpC* VGF::GGR | Expression of *B. subtilis* ClpC VGF::GGR with $His_6$-tag | This work |
| **25** | pET28a c*lpC* VGF::IGF | Expression of *B. subtilis* ClpC VGF::IGF with $His_6$-tag | This work |
| **26** | pET28a *clpC* E280A E618A | Expression of *B. subtilis* ClpC E280A E618A with $His_6$-tag | This work |
| **27** | pET28a *ywlE C7S* | Expression of *B. subtilis* YwlE C7Swith His6-tag | This work |
| **28** | pMAD *clpC* VGF::IGF | For *B. subtilis* markerless point mutation *clpC* V671I (VGF::IGF) *in cis*, *ery* | This work |

**Table S3: List of primers**

| **Nr.** | **Plasmid** | **Primer** | **Sequence** |
|---|---|---|---|
| 1 | 6 7 | SalI_RBS_ClpC_for_IH | GCGTCGACAGAGAACAAGGAGGGGCTACAAATGATGTTTGGAAGATTTACAG |

| | | | |
|---|---|---|---|
| 2 | 6 7 | SphI_ClpC_rev_IH | GCGCATGCTTAATTCGTTTTAGCAGTCG |
| 3 | 8 11 12 13 | BsmBI_McsB_for_Hind III | CCAGTGCGTCTCAAGCTTAAGGAGGGGCTACAAATGTCGCTAAAGCATTTTATTCAGG |
| 4 | 8 12 | SphI_McsB_rev_IH | GCGCATGCTCATATCGATTCATCCTCCTGTC |
| 5 | 9 | SalI_MecA_for_IH | GCGTCGACAGAGAACAAGGAGGGTACAAATGGAAATTGAAAGAATTAACG |
| 6 | 9 | SphI_MecA_rev_IH | GCGCATGCCTATGATGCAAAGTGTTTTTTTATCGTTTCTAGAGCGTGCTCTGAAAT |
| 7 | 10 | SalI_YpbH_rev_IH | GCGTCGACAGAGAACAAGGAGGGTACAAATGCGGCTTGAGCGTCTGAA |
| 8 | 10 | SphI_YpbH_rev_IH | GCGCATGCTTATGAAAAATGAGTTTGTATCG |
| 9 | 11 | SphI_GFP_rev_IH | GCGCATGCTCATTATTTGTATAGTTCATCCATGC |
| 10 | 12 13 16 | McsB_Link_for_IH | AAAAGACAGGAGGATGAATCGATAGCAAGTGGAATGAGTAAAGGAGAAGAACTTTTC |
| 11 | 12 13 16 | McsB_Link_rev_IH | GAAAAGTTCTTCTCCTTTACTCATTCCACTTGCTATCGATTCATCCTCCTGTCTTTT |
| 12 | 16 | BamHI_McsB_for_IH | CCCCGGATCCATGTCGCTAAAGCATTTTATTC |
| 13 | 16 | NcoI_McsB_rev_IH | CCCCCCATGGTCATATCGATTCATCCTCCTGTC |
| 14 | 14 | YwlE_SalI_for_IH | GCGTCGACAGAGAACAAGGAGGGTACAAATGGATATTATTTTTGTCTGTACTGG |
| 15 | 14 15 | YwlE_SphI_rev_IH | GCGCATGCTCATTATCTACGGTCTTTTTTCAGC |
| 16 | 15 | YwlE_C7S_SalI_for_IH | GCGTCGACAGAGAACAAGGAGGGTACAAATGGATATTATTTTTGTCTCTACTGGAAATAC |
| 17 | Construction of the *ypbH::cat* mutant | YpbH_mut1_for_IH | CCCACTTGGTTATAATCGGCGG |
| 18 | | YpbH_mut2_rev_IH | ACTTAAGGGTAACTAGCCTCGCCGAACGTTCCCTCCTGCCTCTACTTGC |
| 19 | | YpbH_mut3_cm_for_IH | TCAAATTTAAGGAGAATCTCATCGGCGAGGCTAGTTACCCTTAAGT |
| 20 | | YpbH_mut4_cm_rev_IH | AAATCAGCAGTTTTCCGTTGCATTCCAATAGTTACCCTTATTATCAAG |
| 21 | | YpbH_mut5_for_IH | CTTGATAATAAGGGTAACTATTGCGATACAAACTCATTTTTCATAA |
| 22 | | YpbH_mut6_rev_IH | AATTCTTGAATACTCATCCATCATCC |

| | | | |
|---|---|---|---|
| 23 | 17 24 25 26 | NcoI_ClpC_for_RK | CATGCCATGGGGTTTGGAAGATTTACAGA |
| 24 | 17 24 25 26 | XhoI_ClpC_rev_RK | CCGCTCGAGATTCGTTTTAGCAGTCGTT |
| 25 | 21 | NcoI_MecA_for_RK | CATGCCATGGAAATTGAAAGAATTAACGAGC |
| 26 | 22 | BsaI_YwlE_for_RK | GGTCTCCCATGGATATTATTTTTGTCTG |
| 27 | 27 | BsaI_YwlE_C7S_for_RK | GGTCTCCCATGGATATTATTTTTGTCAGCACTGGAAATACGTGC |
| 28 | 22 27 | XhoI_YwlE_rev_RK | CCGCTCGAGTCTACGGTCTTTTTTCAGCTG |
| 29 | 23 | NcoI_ClpP_for_RK | CCATGGGTTTAATACCTACAGTCATTGAAC |
| 30 | 23 | XhoI_ClpP_rev_RK | CCGCTCGAGCTTTTTGTCTTCTGTGTGA |
| 31 | 25, 28 | ClpC_VGF::IGF_for_RK | GGAGCAAGTGAGCTAAAACGCAATAAATATATTGGCTTTAACGTTCAGGAT |
| 32 | 25, 28 | ClpC_VGF::IGF_rev_RK | GTCTTTATGATTTTGAGTTTCATCCTGAACGTTAAAGCCAATATATTTATTGCG |
| 33 | 28 | BamHI_ClpC_for_NM | CCCCGGATCCATGATGTTTGGAAGATTTACAG |
| 34 | 28 | NcoI_ClpC_rev_NM | CCCCCCATGGTTAATTCGTTTTAGCAGTCG |

**Table S4: Peptides used for YwlE-His quantification**

| Peptide sequence (*indicates labelled amino acid) | Stable-isotope labelled amino acid | mass endogenous peptide $[M+H]^{1+}$ | Mass synthetic peptide |
|---|---|---|---|
| ATPHAVEALFEK* | $^{13}C_6{}^{15}N_2$-Lysine | 1312.68958 | 1320.70378 |
| QIIASQFGR* | $^{13}C_6{}^{15}N_4$-Arginine | 1019.56325 | 1023.57152 |
| SAGVFASPNGK* | $^{13}C_6{}^{15}N_2$-Lysine | 1034.52873 | 1042.54073 |
| QTRDELEELLR* | $^{13}C_6{}^{15}N_4$-Arginine | 1401.73323 | 1411.7415 |
